# Pithoviruses are invaded by repeats that contribute to their evolution and divergence from cedratviruses

**DOI:** 10.1101/2023.03.08.530996

**Authors:** Sofia Rigou, Alain Schmitt, Jean-Marie Alempic, Audrey Lartigue, Peter Vendloczki, Chantal Abergel, Jean-Michel Claverie, Matthieu Legendre

## Abstract

*Pithoviridae* are amoeba-infecting giant viruses possessing the largest viral particles known so far. Since the discovery of *Pithovirus sibericum*, recovered from a 30,000-y-old permafrost sample, other pithoviruses, and related cedratviruses, were isolated from various terrestrial and aquatic samples. Here we report the isolation and genome sequencing of two *Pithoviridae* from soil samples, in addition to three other recent isolates. Using the 12 available genome sequences, we conducted a thorough comparative genomics study of the *Pithoviridae* family to decipher the organization and evolution of their genomes. Our study reveals a non-uniform genome organization in two main regions: one concentrating core genes, and another gene duplications. We also found that *Pithoviridae* genomes are more conservative than other families of giant viruses, with a low and stable proportion (5% to 7%) of genes originating from horizontal transfers. Genome size variation within the family is mainly due to variations in gene duplication rates (from 14% to 28%) and massive invasion by inverted repeats. While these repeated elements are absent from cedratviruses, repeat-rich regions cover as much as a quarter of the pithoviruses genomes. These regions, identified using a dedicated pipeline, are hotspots of mutations, gene capture events and genomic rearrangements, that contribute to their evolution.

## Introduction

*Pithoviridae* are amoeba-infecting giant viruses possessing the largest known viral particles. The prototype of the family, *Pithovirus sibericum*, was recovered almost 10 years ago from a 30’000-y-old permafrost sample (Legendre et al. 2014). Following this discovery, 6 additional isolates, all infecting *Acanthamoeba castellanii*, have been sequenced (Andreani et al. 2016; Levasseur et al. 2016; Bertelli et al. 2017; Rodrigues et al. 2018; Jeudy et al. 2020). Their dsDNA circular genomes range from 460 to 686 kb. The *Pithoviridae* are composed of two main clades: the pithoviruses and the cedratviruses. Both possess ovoid-shaped virions, capped by a cork-like structure at one extremity for the former and at both extremities for the latter.

*Pithoviridae* have mostly been isolated from permafrost (Legendre et al. 2014; Jeudy et al. 2020; Alempic et al. 2023) and sewage samples (Levasseur et al. 2016; dos Santos Silva et al. 2018). Metagenomic surveys have also revealed *Pithoviridae*-like sequences in deep-sea sediments (Bäckström et al. 2019), in forest soil samples (Schulz et al. 2018), and their high abundance in permafrost (Rigou et al. 2022). In every case, a phylogeny of the metagenomic viral sequences showed that they branch outside the clade of isolated *Pithoviridae*, suggesting that new viral species are yet to be discovered (Rigou et al. 2022).

Genomic gigantism has been observed several times in the virosphere, among viruses infecting prokaryotes, such as “huge” (Al-Shayeb et al. 2020) and “jumbo” phages (Yuan and Gao 2017), or eukaryotes, as in the *Nucleocytoviricota* phylum to which the *Pithoviridae* family belongs. But its origin remains a mystery as most giant viruses’ genes have no known origin. Furthermore, *Pithoviridae* and their relatives are good models to study viral gigantism as there is a variety of genome (and virion) size among the viral order they belong to: the *Pimascovirales* (Lefkowitz et al. 2018; Koonin et al. 2020). The latter is formed by *Iridoviridae*, *Ascoviridae* and *Marseilleviridae* on one side, and *Pithoviridae*, *Orpheovirus*, and related viruses known from metagenomics such as *Hydrivirus* (Rigou et al. 2022), on the other side. The closest isolated relative to *Pithoviridae* is *Orpheovirus*, with a much larger, 1.6 Mb, genome (Andreani et al. 2018). *Orpheovirus* infects *Vermamoeba vermiformis*, while *Pithoviridae* and *Marseilleviridae* both infect *Acanthamoeba*. Some authors consider *Orpheovirus* to be part of *Pithoviridae* (Aylward et al. 2021), although we chose not to, considering the few genes they share (Andreani et al. 2018; Queiroz et al. 2023). *Hydrivirus* also has a 1.5 Mb genome, in contrast with *Pithoviridae* and other *Pimascovirales* such as *Marseilleviridae* with only 350 kb genomes. In *Nucleocytoviricota*, massive horizontal gene transfers from their host (Moreira and Brochier-Armanet 2008) and gene duplications (Filée and Chandler 2008) have been proposed as the driving force behind their expanded genome size. Another mechanism proposed in *Pandoraviridae* is *de novo* gene creation from intergenic regions (Legendre et al. 2018). Whatever the main evolutionary process at play, different families of giant viruses exhibit inhomogeneity in their genomes, by having a “creative” part and a “conservative” one. This pattern is revealed by an unequal distribution of core genes, duplicated genes and genomic rearrangements, preferentially concentrated in one half of the genome (Legendre et al. 2018; Blanca et al. 2020; Christo-Foroux et al. 2020).

Another factor that might shape giant viruses’ genomes are transposons. For instance different *Pandoraviridae* are known to harbor Miniature Inverted Transposable Elements (MITEs) (Zhang et al. 2018). These non-autonomous class II transposable elements are composed of terminal inverted repeats separated by an internal sequence that lacks the transposase gene. Thus, they rely on an autonomous transposon for transposition (Zhang et al. 2001). Their target sites are often as simple as AT dinucleotides that give rise to target site duplication (TSD) (Ge et al. 2017). In *Pandoravirus salinus*, the transposon probably associated to these MITEs has been found in the genome of the *A. castellanii* cellular host (Sun et al. 2015). The *Pithovirus sibericum* genome also contains many copies of a 140-nucleotides-long palindromic repeated sequence in non-coding regions (Legendre et al. 2014). The nature of these repeated sequences, also found in *Pithovirus massiliensis* (Levasseur et al. 2016) remains unknown. Surprisingly, cedratviruses are completely devoid of such sequences (Andreani et al. 2016).

In this study, we report the genome sequences of two new *Pithoviridae* viruses isolated from soil samples (cedratvirus borely and cedratvirus plubellavi), in addition to the recently isolated cedratvirus lena (strain DY0), cedratvirus duvanny (strain DY1) and pithovirus mammoth (strain Yana14) (Alempic et al. 2023). The comparative analysis of these sequenced genomes, complemented with previously published *Pithoviridae* sequences (Legendre et al. 2014; Levasseur et al. 2016; Bertelli et al. 2017; Rodrigues et al. 2018; Jeudy et al. 2020), provides insight into the gene distribution and the evolution of the family. In addition, an in-depth study of pithoviruses’ genomes reveals that they are highly structured in regions composed of two main inverted repeats, that have massively colonized their genomes and influenced their evolution.

## Results

### *Pithoviridae* isolation from soil samples and genome sequencing

We isolated two new viruses that belong to two species of cedratviruses (*Cedratvirus borely* and *Cedratvirus plubellavi*), both infecting *A. castellanii*, from two soil samples located 10m away in a French park (43°15’34.0″N, 5°22’58.9″E and 43°15’34.3″N, 5°22’59.2″E, respectively). As shown for cedratvirus plubellavi in Fig. S1, they possess a typical lemon-like *Cedratvirus* morphology with two corks, one at each apex of the particle. We next sequenced their genomes. In addition, we assembled and annotated the ones of three recently reported *Pithoviridae* isolated from various Siberian environments (Alempic et al. 2023), including a *Pithovirus* from frozen soil containing mammoth wool (*Pithovirus mammoth*), a *Cedratvirus* from the Lena river in Yakoutsk (*Cedratvirus lena*) and another *Cedratvirus* (*Cedratvirus duvanny*) from a melting ice wedge in the Duvanny yar permafrost exposure (Table 1). Long-read sequences were available for three viruses. They turned out to be essential for the completeness of the *Pithovirus mammoth* assembly, while they only had a minor effect on the *Cedratvirus borely* assembly and no effect at all on the *Cedratvirus plubellavi* one (Table S1). The three genomes were successfully circularized, as shown by the homogeneous long read coverage along the genomes artificially linearized at 4 equidistant positions (Fig. S2). It should be noted that the circularity of the *Pithoviridae* genomes has previously been proven by a Pulse-Field Gel Electrophoresis experiment on *Cedratvirus kamchatka* DNA (Jeudy et al. 2020).

**Table 1.** Genome metrics of sequenced *Pithoviridae* from this study compared to previously published isolates. The names of the *Pithoviridae* sequenced in this study are underlined while the names in bold represent the mean and standard deviation of the group considering all isolates. *Cedratvirus* clades follow the ones defined in (Jeudy et al. 2020) and are shown in Figure 1.

|  | Real genome |  |  | Without repeats |  |  |
| --- | --- | --- | --- | --- | --- | --- |
|  | Length (kb) | GC% | Coding density | Length (kb) | GC% | Coding density |
| <u>*Pithovirus mammoth*</u> | 610 | 35.8 | 0.7 | 469 | 39.5 | 0.9 |
| **<u>pithoviruses</u>** | 637 ± 40.15 | 35.6 ± 0.13 | 0.6 ± 0.03 | 485 ± 0.04 | 39.5 ± 0.04 | 0.9 ± 0.02 |
| <u>*Cedratvirus borely*</u> | 570 | 42.8 | 0.8 | 553 | 42.8 | 0.8 |
| <u>*Cedratvirus plubellavi*</u> | 568 | 42.8 | 0.8 | 552 | 42.8 | 0.9 |
| **<u>cedratviruses clade A</u>** | 573 ± 10.49 | 42.8 ± 0.02 | 0.8 ± 0.01 | 556 ± 0.01 | 42.8 ± 0.01 | 0.8 ± 0.01 |
| <u>*Cedratvirus lena*</u> | 466 | 40.8 | 0.8 | 434 | 40.7 | 0.9 |
| <u>*Cedratvirus duvanny*</u> | 472 | 40.8 | 0.8 | 440 | 40.8 | 0.9 |
| **<u>cedratviruses clade B</u>** | 468 ± 3.5 | 40.7 ± 0.1 | 0.8 ± 0.02 | 441 | 40.7 ± 0.09 | 0.9 ± 0.01 |
| **<u>cedratviruses clade C</u>** | 460 | 43 | 0.8 | 445 | 42.9 | 0.9 |

All included, 12 *Pithoviridae* genome sequences are now available (Table S2) for a comparative study of the family.

### Pithoviridae phylogeny

To get insight into the *Pithoviridae* family evolution, we next performed a phylogenetic reconstruction of the 12 genomes in addition to the more distantly-related *Orpheovirus* (Andreani et al. 2018) and *Hydrivirus*, the only complete *Pithoviridae*-like genome assembled from metagenomics data (Rigou et al. 2022). As shown in Figure 1, *Orpheovirus* and *Hydrivirus* are the most divergent, pithoviruses and cedratviruses split into two well established clades, and cedratviruses can be further divided into 3 previously defined clades (Jeudy et al., 2020). Although *Hydrivirus* and *Orpheovirus* cluster in a well-supported clade, they diverge from each other (Average Amino-acid Identity, AAI = 31%) more than cedratviruses from pithoviruses (AAI = 42.2% ± 0.2). In addition, *Hydrivirus* and *Orpheovirus* only share 140 HOGs (Hierarchical Orthologous Groups, see Methods), as compared to the more than 1400 genes identified in their respective genomes. This suggests that the group will likely split into better defined clades as new related viruses are added.

**Figure 1.**
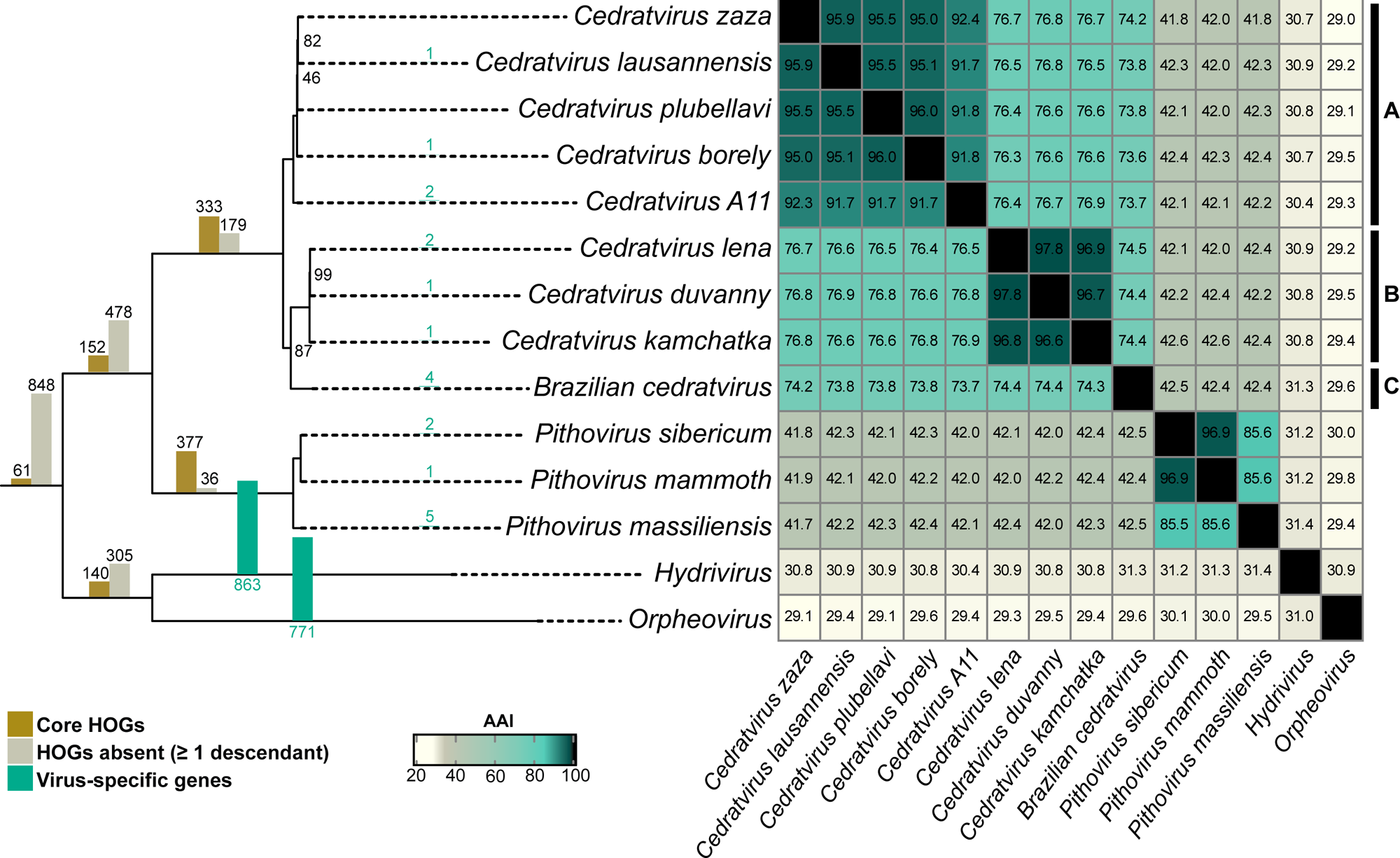
Phylogeny and average amino-acid identity of the *Pithoviridae* and their closest relatives. The phylogeny (left) was built from the concatenation of shared single copy HOGs (Hierarchical Orthologous Groups) applying the LG+F+G4 evolutionary model. Bootstrap values are indicated or are 100% otherwise. The bars on each branch represent the number of shared HOGs and other HOGs that were recomputed by OrthoFinder according to this tree. The heatmap (right) shows the average amino-acid identity (AAI) between viruses. The right-most bars (labeled A, B and C) indicate previously determined *Cedratvirus* clades (Jeudy et al. 2020).

Consistent with the phylogeny, the codon usage pattern shows a similar trend, with cedratviruses tightly clustered together, as for pithoviruses, and *Orpheovirus* being the most distant (Fig. S3). This is in line with the fact that the *Pithoviridae* and *Orpheovirus* infect different laboratory hosts (Andreani et al. 2018).

Within cedratviruses or pithoviruses, genomes are globally collinear despite several rearrangements (Fig. S4). *Pithovirus massiliensis* shows one major inversion and one translocation compared to the two other pithoviruses. Both *Cedratvirus kamchatka* and *Brazilian cedratvirus* exhibit many rearrangements compared to clade A.

### Heterogeneity within the genomes of *Pithoviridae*

The comparative genomics studies of other giant virus families previously highlighted a biased evolution of their genomes with a “creative” and a “conservative” part (Legendre et al. 2018; Blanca et al. 2020). We thus looked for a similar trend in the *Pithoviridae* genomes. As shown in Figure 2A, core genes are not uniformly distributed along the artificially linearized pithoviruses’ genomes, with a high concentration at one half containing the ATP-dependent DNA ligase. Likewise, core genes are also very scarce in the other half of the cedratviruses genomes. This pattern contrasts with gene duplications that seem to occur in specific hotspots preferentially located with the accessory genes (Fig. 2B). Altogether, this data shows a shared non-uniform architecture of the *Pithoviridae* genomes.

**Figure 2.**
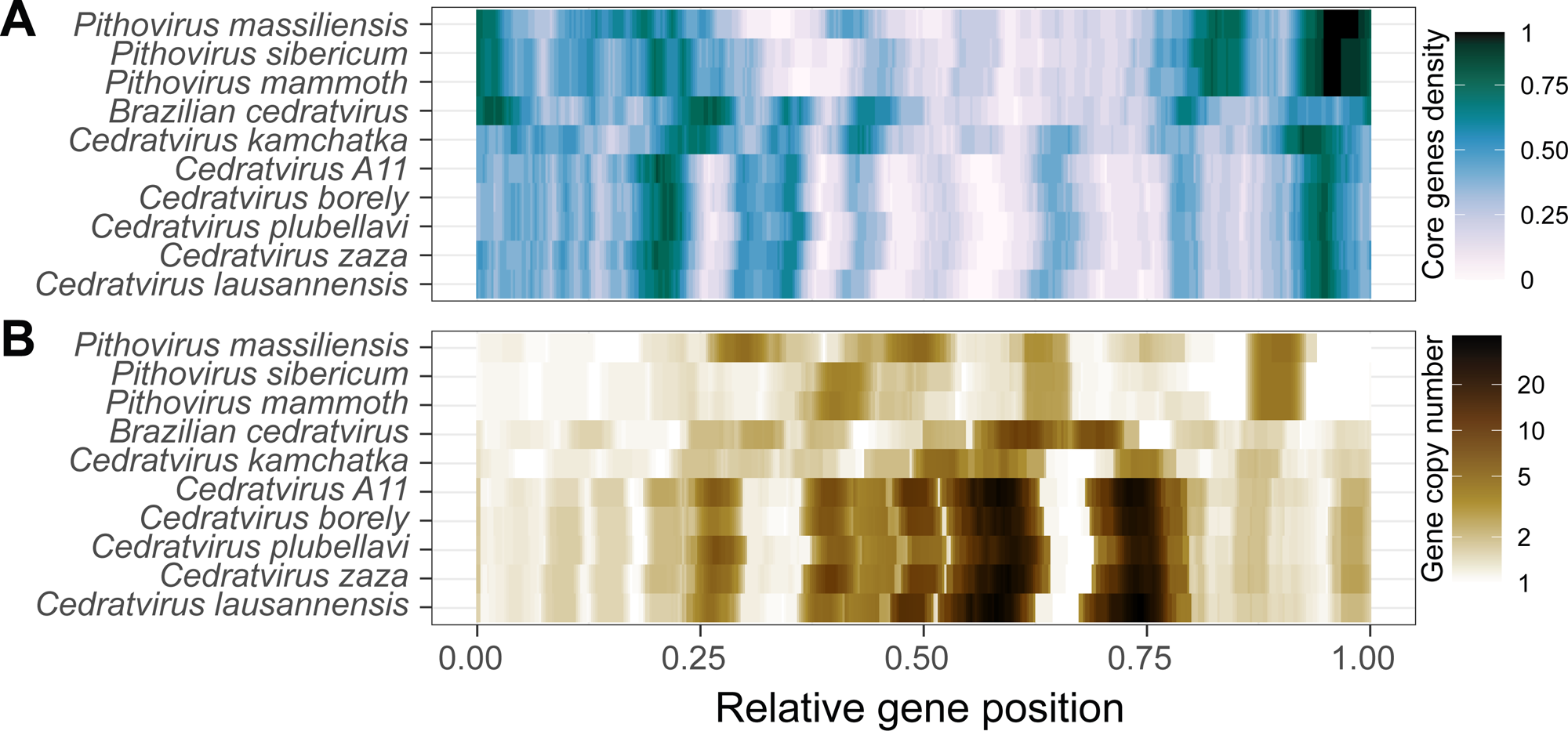
Non-uniform distribution of core and duplicated genes. (A) Density of core genes within a sliding window of 21 ORFs. (B) Average gene copy number within the HOGs containing each of the genes of the sliding window.

We next questioned whether *Pithoviridae* DNA was differently epigenetically modified between the “creative” and the “conservative” regions. We analyzed the PacBio data previously generated for *Pithovirus sibericum* and *Cedratvirus kamchatka* (Jeudy et al. 2020) and extracted all positions with interpulse duration (IPD) ratios > 4.5 as potentially modified. We found no significant difference with 0.53 modified bases per kb in the “conservative” region and 0.61 in the “creative” region of *Cedratvirus Kamchatka* (chi² test *P*value = 0.34). At this IPD threshold, no *Pithovirus sibericum* nucleotide is predicted to be modified, as previously noticed (Jeudy et al. 2020).

### *Pithoviridae* are conservative compared to other *Nucleocytoviricota*

We next quantified the *Pithoviridae* core- and pan-genomes and compared them to other viral families. The core-genome of cedratviruses is made of 333 ORFs (Open Reading Frames) over 100 amino-acids (Fig. 3A-B) while the one of the whole *Pithoviridae* family is twice as small with an asymptote at 152 ORFs (Fig. 3B).

**Figure 3.**
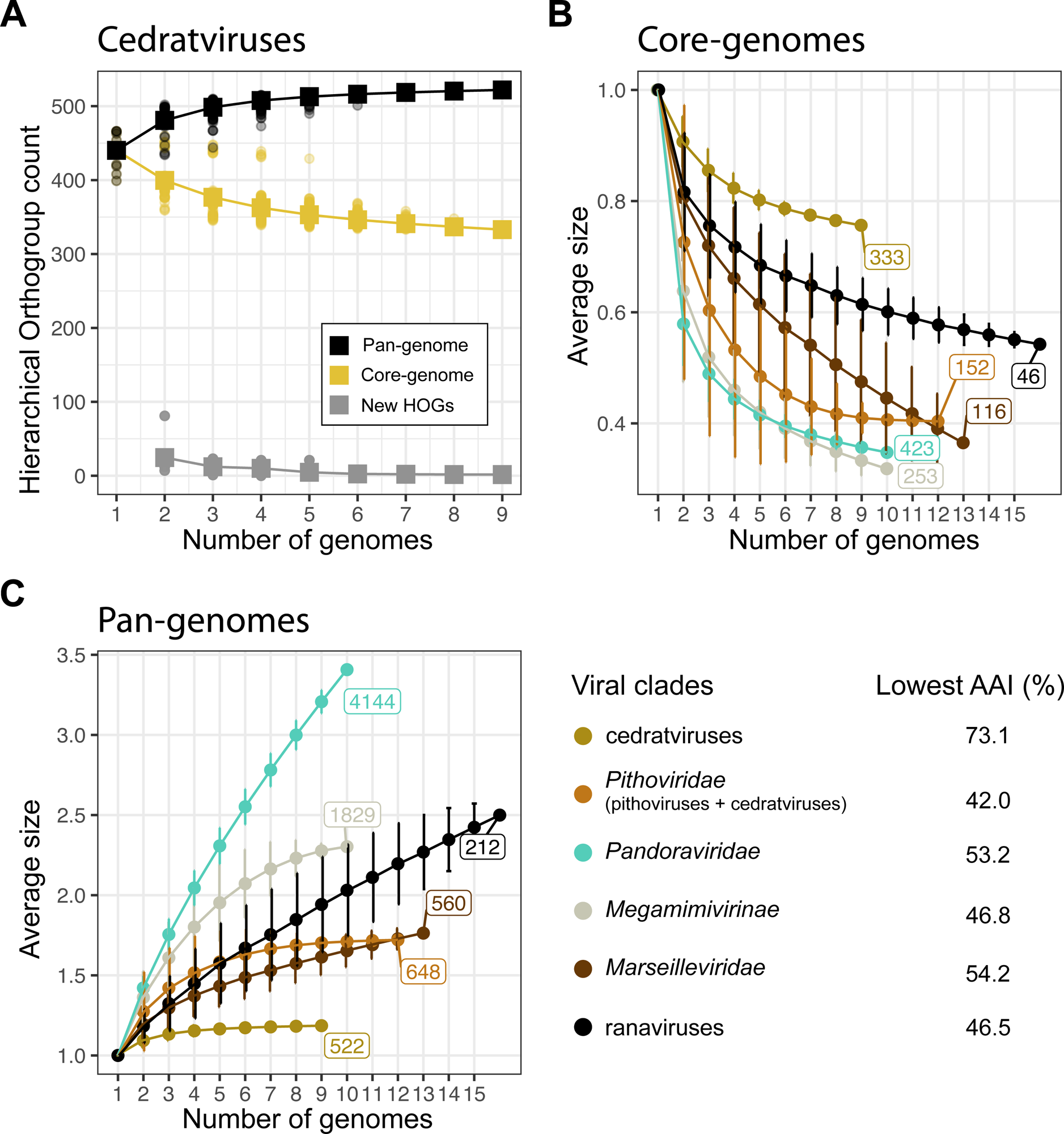
The core and pan-genomes of *Pithoviridae* and other *Nucleocytoviricota*. (A) Pan-genome, core-genome and new HOGs have been estimated for cedratviruses by adding new genomes to a set of previously sequenced genomes in an iterative way (Tettelin et al. 2005). For comparison, the core-genome (B) and pan-genome (C) sizes of other *Nucleocytoviricota* have been estimated in the same iterative way. The pan-genome and core-genome sizes are defined as the relative size in comparison to their initial mean size. The lowest AAI shown in the legend indicates the AAI of the most distant viruses within the set of genomes used for this analysis.

The pan-genomes of cedratviruses and *Pithoviridae* (including cedratviruses and pithoviruses) have both reached a plateau (Fig. 3C), suggesting a so-called “closed” pan-genome. In agreement with this, each new genome brings less than two new HOGs to the cedratviruses (Fig. 3A). The closedness of the cedratviruses and *Pithoviridae* pan-genomes was confirmed by Heaps law models with α estimates of 2 in both cases. Pan-genomes with α < 1 are open and α > 1 closed. By contrast, *Pandoraviridae* and ranaviruses have open pan-genomes with both α estimates of 0.53. Finally, *Megamimivirinae* and *Marseilleviridae* exhibit a closer pan-genome with α estimates of 1.19 and 1.13, respectively. In other words, *Pithoviridae* appear to be much more conservative (*i.e.* closer pan-genome) than other *Nucleocytoviricota* (Fig. 3C), suggesting that, unlike for *Pandoraviridae,* continuous *de novo* gene creation might not be a significant process in their evolution (Legendre et al. 2018).

It is worth noting, however, that the phylogenetic breadth of each group has a direct impact on the pan-genome and core genome trends. According to the lowest AAI within each group (Fig. 3), cedratviruses are more closely related than the other groups (lowest AAI = 73%) whereas *Pithoviridae* contains distant viruses with a lowest AAI value of 42 % (Fig 3). Both phylogenetic groups display a closed pan-genome. In concordance with this apparent conservative evolution, cedratviruses and pithoviruses specific genes are mostly shared within their respective genomes, in contrast to *Pandoraviridae* and *Marseilleviridae* that exhibit a much larger fraction of accessory genes within their sub-clades (Fig. S5). A better sampling of the two *Pithoviridae* clades (cedratviruses and pithoviruses) will be needed to confirm the closedness of the family pan-genome.

### Gene duplication and HGT in *Pithoviridae*

Next, we investigated gene duplication as a possible important cause of viral genome gigantism (Filée and Chandler 2008). Gene duplications occurred all along the history of *Pithoviridae*, even during the short divergence time separating the closely related *Pithovirus sibericum* and *Pithovirus mammoth*. They mostly occurred in the vicinity of their original copy with a median distance of 6872 bp in cedratviruses and 1575 bp in pithoviruses. Overall, from 14 % to 28 % (median = 19 %) of the *Pithoviridae* genes come from a duplication event (Fig. 4), in line with other *Nucleocytoviricota* such as *Marseilleviridae* (16 %), *Pandoraviridae* (15 %) and *Megamimivirinae* (14 %). Within cedratviruses, gene duplications largely explain genome size variations between clade A and clades B-C, with 27.4 ± 0.9 % in clade A and 18.5 ± 1.7 % in clades B-C (Fig. 4). Consistently, the most duplicated gene, coding for an ankyrin-repeat protein, is present in 50 copies in clade A cedratviruses and only 20 copies in clades B-C. Likewise, the related *Orpheovirus* and *Hydrivirus* very large genomes exhibit high rates of gene duplications, with 42% and 27% respectively (Fig. 4). By contrast, there is a converse pattern in-between pithoviruses and cedratviruses, the latter, displaying higher duplication rates while having smaller genomes, suggesting that another factor is at play.

**Figure 4.**
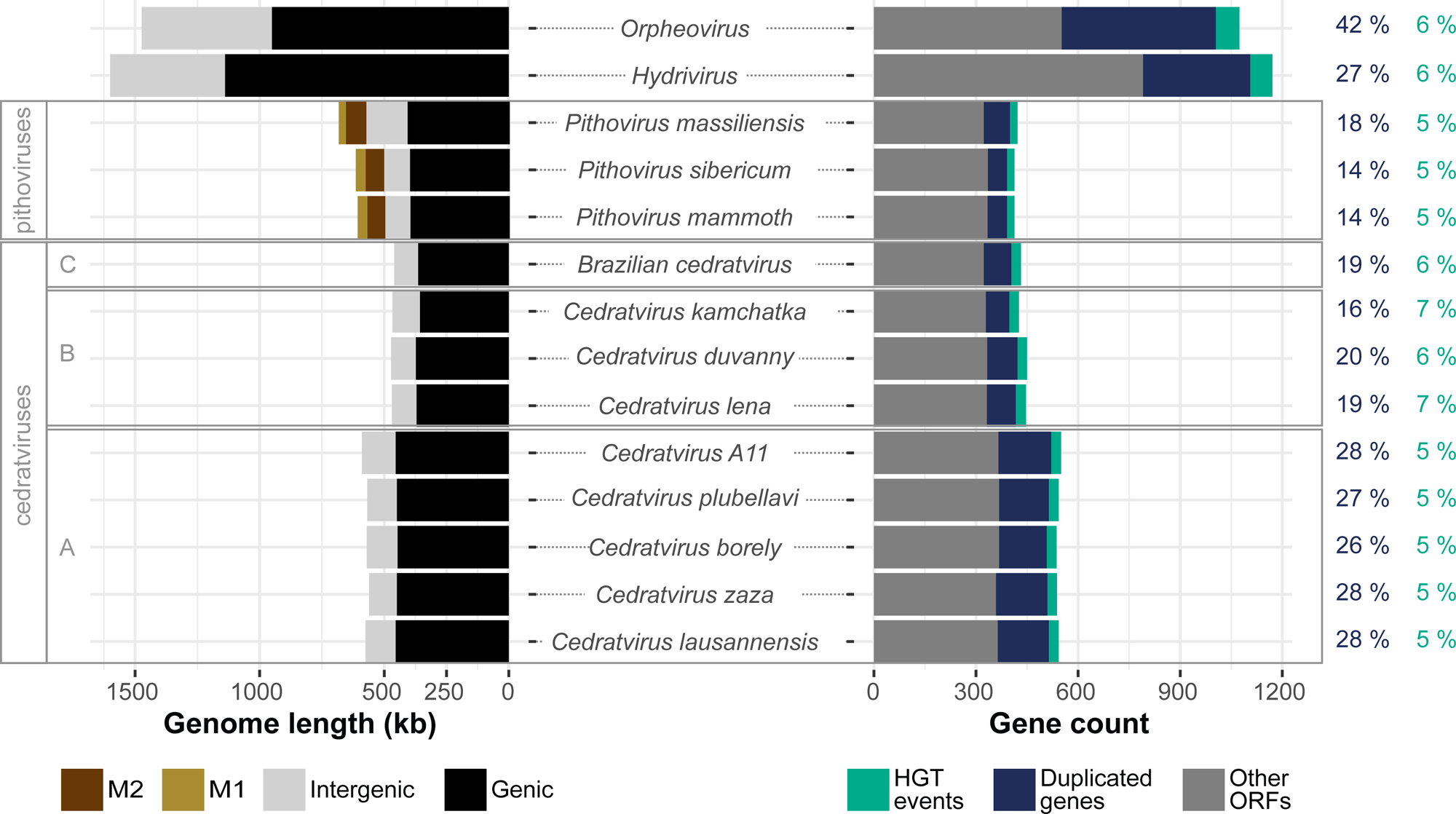
Genome and gene content statistics of *Pithoviridae* and relatives. The left panel presents the nucleotide content of the different genomes with clades labels on the left (see Fig. 1). M1 and M2 correspond to inverted repeats (see further). The right panel shows their composition in ORFs. The percentage of genes that arose from a duplication event (in blue) and the percentage of HGT events toward each genome (in green) are shown on the right.

We next investigated horizontal gene transfers (HGTs) towards our viruses based on the HOG phylogenetic trees complemented with homologous sequences (see Methods), as a possible source of genome size increase. It turned out that HGTs are far less frequent than gene duplications with a stable fraction of 5% to 7% of the gene content across *Pithoviridae* and in *Orpheovirus* and *Hydrivirus* (Fig. 4).

The largest proportion of *Pithoviridae* HGTs come from eukaryotes (42 ± 2 %) closely followed by those originating from Bacteria (41 ± 3 %), (Fig. S6). The HGT from Eukaryota do not clearly point to known hosts. Most often, the root of the HGT is ancient, branching before or in-between Discosea and Evosea, two classes of amoebas (Fig. S6). We also estimate that 10 % of the HGT events came from another virus.

Overall, the low rate of HGT in *Pithoviridae* is consistent with the closedness of their pan-genome and thus cannot account for the difference in genome sizes between cedratviruses and pithoviruses, hinting again at a different factor.

### Two types of inverted repeats have massively colonized the genomes of pithoviruses

Repeat content is another factor that could strongly influence genome size. Indeed, it has been shown that pithoviruses genomes are shaped by intergenic interspersed palindromic repeat sequences (Legendre et al. 2014). These are present in clusters and usually separated by 140 nucleotides (median). After masking these sequences (Fig. 5A-B) from the genomes, we identified additional repeats close to the masked regions (Fig. 5C-D). By running the MUST (Ge et al. 2017) and MITE-Tracker (Crescente et al. 2018) tools, we found that both types of repeats were identified as putative MITEs that we referred to as M1 and M2.

**Figure 5.**
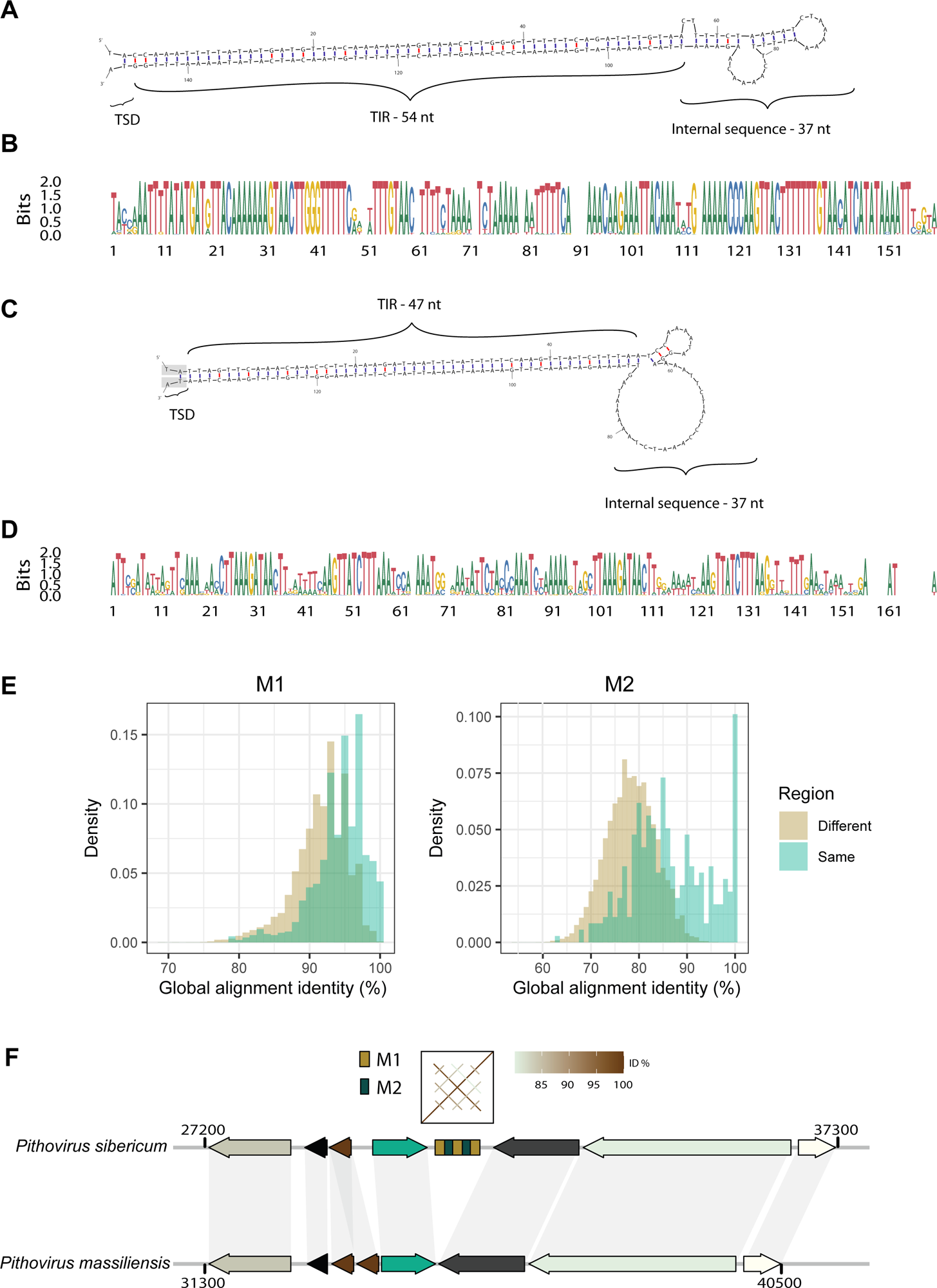
Main repeats found in pithoviruses. DNA folding structures of the reference sequence for M1 (A) and M2 (C) repeat clusters respectively. Their free energy ΔG is of -79.2 and -65.5 kcal/mol. TSD is for Target Site Duplication and TIR for Terminal Inverted Repeat. The TSD highlighted in grey in (C) indicates that the dinucleotide is shared between M1 and M2 when the repeats are next to each other. (B) and (D) are the alignment logos of all the sequences in the clusters of M1 and M2 respectively. (E) Pairwise identity percentage in-between M1 (left) and M2 (right) repeats retrieved from the same (green) and from distinct (brown) regions. The pairwise identity percentages were calculated using the needle tool from the EMBOSS package. Both distributions are significantly different (*P*values < 10^-15^, wilcoxon test). (F) Example of a repeated region present in *Pithovirus sibericum* but absent from *Pithovirus massiliensis* in a syntenic region of their genomes.

We designed a pipeline that define repeat-rich regions by automatically identifying and clustering repeat sequences (see Methods and Fig. S7). The reference M1 and M2 are palindromic with terminal inverted repeats (TIRs) of 54 and 47 bp, respectively, and an internal sequence of 37 bp (Fig. 5A,C). The alignment of the extremities of M1 and M2 (and of repeat-rich regions) suggests TA as a putative target site duplication (Fig. S8). When combined together, the M1 and M2 sequences represent as much as 18.4%, 18.2% and 16.1% of the genomes of *Pithovirus sibericum*, *Pithovirus mammoth* and *Pithovirus massiliensis*, respectively (Fig. 4), and 21 to 24% when all kinds of repeats are considered. It is worth noting that when *Pithovirus sibericum* was first discovered, it was estimated that 21% of its genome was covered by repeats (Legendre et al. 2014). This fraction includes both M1 and M2 repeats, although the latter were not identified at the time. Unlike duplicated and core genes (Fig. 3), repeats are not concentrated in specific genomic regions but are uniformly distributed along the pithoviruses genomes (Kolmogorov-Smirnov test against uniform distribution *P*value = 0.6). Our pipeline also provided an extensive description of the structure of the repeated regions resulting in the following rules:

1. M2 can never be seen in a repeat region without M1
2. M1 can be seen without M2
3. When several M1 are present in a region, they are always separated by a sequence of about 140 bases, whether M2 is present or not
4. When several M2 are present in a region, they are separated by M1

The most common structure of the repeated regions in the three pithoviruses genomes is: (M1-M2){1 to 8 times}-M1. In *Pithovirus sibericum*, M1 is present 515 times and M2, 371 times.

For comparison, we also tested RepeatModeler on the *Pithovirus sibericum* genome and identified four families of repeats containing 240, 121, 56 and 30 sequences, respectively, covering 25.4% of the genome. As shown by a dotplot of the family consensus sequences (Fig. S9), they partially overlap each other and contain the basic units found by our procedure (M1 and M2). We thus pursued the pithoviruses genomes analyses with the core-units of the repeats identified by our method.

*Pithovirus mammoth* has a very similar number of regions containing M1 and M2 (Table S3) but the number of M1 or M2 copies per orthologous region is often different. Thus, the differences most often correspond to the extension or contraction of existing repeated regions rather than insertions in a repeat-free region. The extension of existing repeat regions is supported by the fact that repeats from the same region are more similar to each other than repeats from different regions (Fig. 5E, *P*values < 10).

Insertion of repeats in repeat-free regions is also necessary to explain the observed high number (> 109) of repeat regions, (Table S3). Insertions and/or excisions have happened several times since the divergence of *Pithovirus sibericum* and *Pithovirus massiliensis*, as exemplified by a repeated region in *Pithovirus sibericum* that is absent from the cognate syntenic orthologous region in *Pithovirus massiliensis* (Fig. 5F). This particular example bares no signs of target site duplication, making transposition mechanism less likely.

### M1 and M2 repeats in metagenomics data

We next questioned whether M1 and M2 repeats were present outside the 3 pithoviruses analyzed in this study by screening the non-redundant NCBI database (that includes the genome of *A. castellanii*) and metagenomic *Nucleocytoviricota* assembled sequences, but none were found. As M1 and M2 sequences might be present in metagenomes but lost during the assembly process, we further looked for reads matching these sequences using the NCBI PebbleScout tool (https://Pebblescout.ncbi.nlm.nih.gov/). We found 28 metagenomic datasets with reads matching M1 or M2 with a PebbleScout score > 70 (Table S4). These correspond to 19 to 1725 reads and 6 to 368 reads matching M1 and M2, respectively, with a BLASTN *E*value<10^-10^. Most of the metagenomic samples correspond to environments for which *Pithoviridae* were previously isolated, namely soil, groundwater and sediments samples (Table S4). We then *de novo* assembled the 10 datasets with highest density of reads matching M1 or M2 and obtained 363 contigs matching those sequences (BLASTN *E*value<10^-10^) among the 16,787,096 assembled contigs. All of them were small, ranging from 211 to 1091 nt and devoid of ORFs. We also searched for the *Pithovirus sibericum* divergent MCP gene (pv_460) in the assembled metagenomic contigs using TBLASTN and found a highly significant match (*E*value<10^-22^) in 9 out of the 10 analyzed datasets. This suggests that the metagenomic contigs matching M1 and M2 might originate from *Pithoviridae*, although we cannot exclude that they belong to other organisms that coexist in the samples.

Furthermore, we found a few *Pithoviridae*-like genomes from metagenomic data that were highly structured by direct repeats (Fig. S10A-B). These constitute 13% of the LCPAC302 pithovirus-like partial genome sequenced from deep-sea sediments (Bäckström et al. 2019). But those repeats have no sequence similarity to M1 or M2. Overall, *Pithoviridae* and *Pithoviridae*-like genomes are highly diverse in repeat content, ranging from none to almost a quarter of their assembled genomes.

### Functional annotation of genes present in repeat-rich regions of pithoviruses

The M1 and M2 repeats in pithoviruses are palindromic (Fig. 5 A,C), such as the MITEs previously identified in pandoraviruses (Sun et al. 2015), and predicted as potential MITEs by MITE searching algorithms. They are also encompassed by potential TSD (Fig. S8), suggesting that they might, at least originally, have integrated the pithoviruses ancestor genome through transposition. To explore this possibility, we analyzed the functional annotation of the pithoviruses repeat-rich regions, but no transposase could be found in current pithoviruses annotations or in M1/M2 containing metagenomic contigs. Instead, we found GO term enrichment for GTP binding and purine nucleoside/ribonucleoside binding (*P*values for *P. sibericum*=0.043, *P. mammoth*=0.016 and *P. massiliensis*=0.0031).

We next performed a remote homology search from protein structure predictions to identify transposase candidates. Alphafold models were built for all proteins, followed by structural alignments using Foldseek. Among the folds obtained, 5 had their best matches with transposases or integrases with Foldseek probability > 0.5 (Table S5). Interestingly, all are within repeat-rich regions and part of multiple copy HOGs. However, the associated Foldseek *E*values were weak, potentially due to a mild confidence in the Alphafold models (average pLDDT = 62). In addition, these genes are small (67 amino acids on average) and under weak selective constraints, with dN/dS ratios of 0.98 +/- 0.7. This suggests that if these transposase-related genes were indeed involved in M1/M2 transposition, they are probably inactive and undergoing pseudogenization.

We next pursued the strategy of structural homology search to increase *Pithovirus sibericum* functional annotation. This resulted in 37 genes with a better functional annotation (Table S6), of which 9 are located in repeat-rich regions. One of those, pv_445, is a Crossover junction endodeoxyribonuclease RuvC-like, that is absent in cedratviruses. By aligning the pv_445 model with the *Fowlpox virus* structural homolog (Li et al. 2020), we identified the DDE active site and other residues important for DNA binding and cleavage (Fig. S11). One could hypothesize that this protein is involved in homologous recombination as a repeat-expansion factor. Finally, the Foldseek alignments revealed a SbcCD subunit D Nuclease (Table S6) that cleaves DNA hairpin structures. Hairpins are dense in repeats, which may increase the instability of those regions. This gene, in conjunction with the DNA double-strand break repair ATPase (pv_215 in *Pithovirus sibericum*), could possibly also be a part of the machinery helping the spread of M1 and M2 sequences.

### Pithoviruses’ repeat-rich regions are hotspots of genetic variability

As repeats constitute a large proportion of pithoviruses’ genomes, we further investigated the genes located in those regions from an evolutionary perspective. Although HGTs are not abundant in pithoviruses (Fig. 4), they are sightly but significantly enriched in repeats regions: 12.4% within *versus* 4.7% outside (chi² test *P*value = 1.7 x 10^-7^, and individual *P*values of 0.002, 0.007 and 0.001 in *Pithovirus sibericum*, *Pithovirus massiliensis* and *Pithovirus mammoth*, respectively).

We also estimated the ancestry of the genes present within these regions compared to other regions. This was performed considering the last common ancestor of all species within each HOG. From that, we observed a significant trend (Cochran-Armitage test *P*value = 2.6 x 10^-3^) whereby newly acquired genes appeared more frequent than ancestral genes in these regions (Fig. 6A). In other words, repeated regions are more prone to gene novelty.

**Figure 6.**
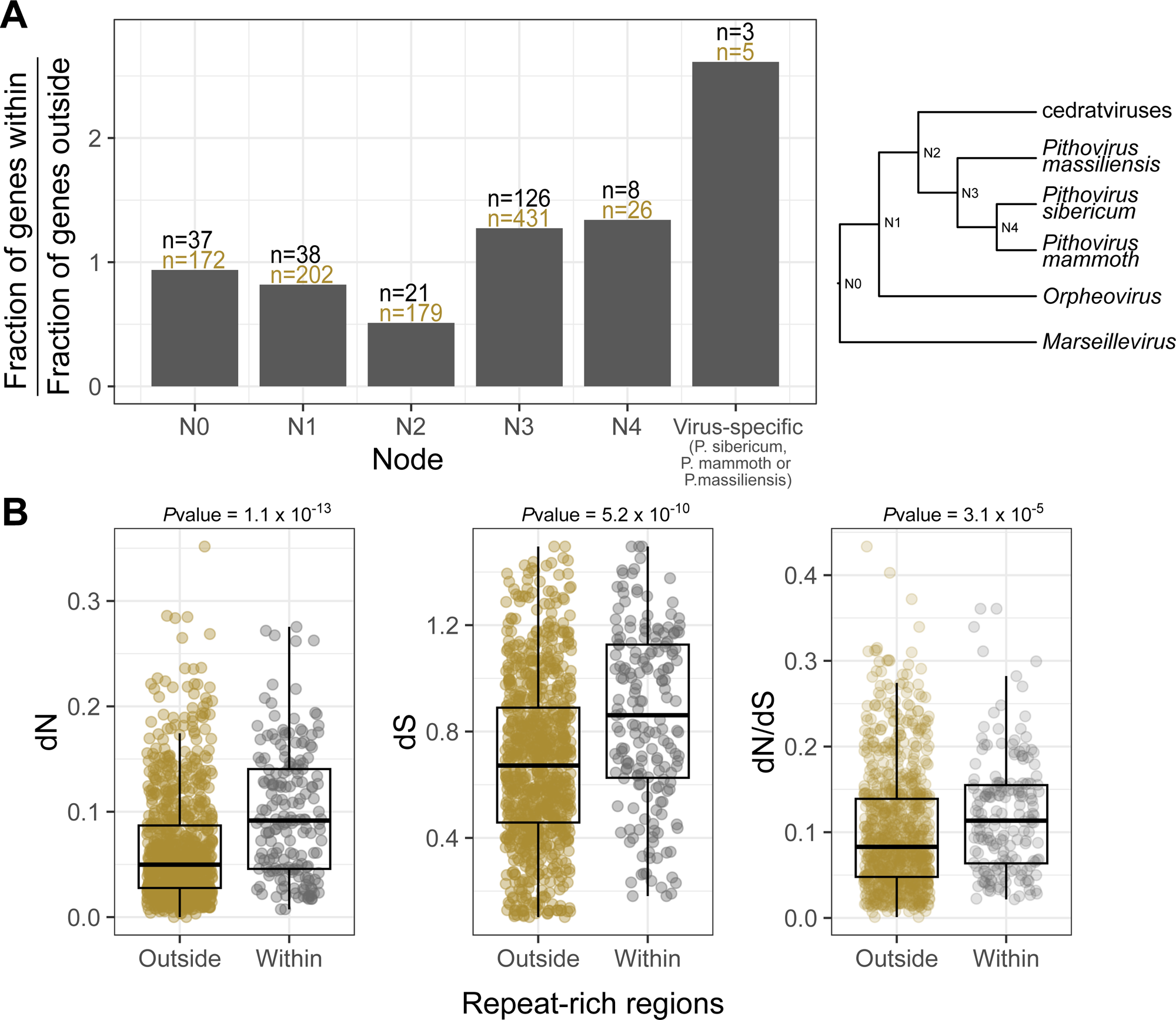
Evolution of ORFs within and outside repeat-rich regions. (A) Ancestry of genes within and outside repeat-rich regions. The ancestry of each gene was estimated considering the last common ancestor of the species present in the cognate HOG. Nodes are ordered from the most ancient to the most recent, as shown in the cladogram next to the plot. (B) dN, dS and dN/dS values for all genes within (gray) or outside (brown) repeat-rich regions detected by our pipeline. *P*values were calculated using Wilcoxon rank tests.

The rates of mutation in repeat-rich *versus* repeat-free regions were compared using orthologous genes. We found that genes located in repeat-rich regions tended to have higher mutation rates for both, synonymous (*P*value = 5.2 x 10^-10^) and non-synonymous (*P*value = 1.1 x 10^-13^) positions (Fig. 6B). All trends are confirmed when considering each individual genome (max *P*value = 7.3 x 10^-4^). In addition, the genes within repeat-rich regions also exhibit higher dN/dS values, thus are less evolutionary constrained (Fig. 6B, *P*value = 3.1 x 10^-5^ and maximum *P*value for individual genomes = 0.027).

Finally, we investigated the frequency of genomic rearrangements located in repeat-rich compared to repeat-free regions. We took advantage of the fact that two pithoviruses (*Pithovirus sibericum* and *Pithovirus mammoth*) were sequenced using long reads and exhibited mostly colinear genomes. We manually inspected orthologous regions of these two viruses to spot potential rearrangement and mutational events. Again, we found that repeat-rich regions were highly enriched in several types of rearrangements compared to repeat-free regions. This includes insertions/deletions, inversions, duplications and substitutions affecting genes, accounting for a total 28 events in repeat-rich regions for only 13 in repeat-free regions (chi^2^ *P*value = 1.42 x 10^-11^, Table S7).

Altogether, these various results establish that pithoviruses repeat-rich regions are hotspots of genetic novelty and undergo relaxed evolutionary constraints.

## Discussion

Here we reported the isolation from soil samples and genome sequencing of two cedratviruses (*Cedratvirus borely* and *Cedratvirus plubellavi*). We also assembled and annotated the genome of *Pithovirus mammoth* recently isolated from 27000-y-old permafrost, of a *Cedratvirus* from fresh water (*Cedratvirus lena*) and another one from melting ice (*Cedratvirus duvanny*) (Alempic et al. 2023). Along with previously described *Pithoviridae*, mostly originating from permafrost (Legendre et al. 2014) and sewage water (Levasseur et al. 2016; Bertelli et al. 2017; dos Santos Silva et al. 2018), these new isolates confirm the ubiquity of this viral family, members of which are present within various aquatic and soil environments. This is also consistent with recent metagenomic surveys exhibiting the presence of *Pithoviridae* in permafrost, forest soils and deep-sea sediments (Bäckström et al. 2019; Rigou et al. 2022).

These 5 additional sequenced strains were combined to 7 previously published genomes to perform a thorough comparative analysis of the *Pithoviridae* family, revealing the organization of their circular genomes. We have found that repeat content is highly diverse within this family. This directly impacts the strategies employed for genome assembly. While addition of long reads only had a limited benefit over short-read only assemblies of cedratviruses, it has a drastic effect on the genome completeness of highly repeated pithoviruses genomes (Table S1). This recently prompted some authors to employ long reads only as a cost-effective strategy for complete genome assembly of giant viruses, including a *Pithovirus* (Hikida et al. 2023).

In all *Pithoviridae* assemblies, we found that their genes are broadly distributed in two distinct regions, one enriched in core genes and the other in gene duplications (Fig. 2). This type of non-uniform genome partition with a “creative” and a conserved region is reminiscent of what has been observed in *Marseilleviridae* (Blanca et al. 2020), a viral family belonging to the same order (*Pimascovirales*) and whose genomes are also circular. However, the two regions are more clearly defined in *Marseilleviridae*, where duplications and accessory genes are evenly dispersed in the “creative” region, while they occur in hotspots in *Pithoviridae* (Fig. 2). Other viral families, that only share a handful of genes (Mönttinen et al. 2021) and have various virion morphology and genome organization (linear or circular), also exhibit this non-uniform distribution of their genes. In *Poxviridae* for instance, core genes are concentrated in the central part of the genome while accessory genes, mostly involved in host-virus interactions, are located at the genome termini (Senkevich et al. 2021). It has been proposed that this accessory partition is a hotspot of frequent gene loss and gain through HGTs (Senkevich et al. 2021), but the few HGT identified in *Pithoviridae* does not support this model. In *Pandoraviridae*, core and essential genes, and those whose proteins are identified in the viral particle, are mostly localized in the left part of the genome, while accessory genes are located on the right part (Legendre et al. 2018; Bisio et al. 2023). This likely reflects ongoing genome increase involving *de novo* gene creation (Legendre et al. 2018) and accelerated gene duplications (Bisio et al. 2023). One could also hypothesize that the partitioning of the genomes is linked to a global epigenetic regulation of gene expression. However, in this work, we found no difference in modified bases densities between the two regions in *Pithoviridae*, which is consistent with the uniform distribution of known methylated motifs in giant viruses’ genomes (Jeudy et al. 2020). Transcriptome analyses of the *Pithoviridae* would be needed to determine whether gene expression profiles are globally different in these large genomic regions. Another factor that might explain the dichotomous partitioning of the *Pithoviridae* genomes is DNA compaction. But while virally-encoded histones have been found in other *Pimascovirales*, namely *Marseilleviridae* (Thomas et al. 2011; Liu et al. 2021), our thorough gene annotation of isolated *Pithoviridae* has not identified such proteins yet. To explore this hypothesis, global 3D structure of the different genomic regions remains to be exanimated by chromosome conformation capture experiments.

Even though *Pithoviridae* genomes are conservative, the cedratviruses and pithoviruses clades exhibit large differences in genome sizes correlated with their repeat contents. The pipeline we designed to study them identified two repeated units in pithoviruses, referred to as M1 and M2. Those repeats share some features with MITEs, such as TIRs (Fig. 5), size and putative TSD (Fig. S8). They are frequently organized in (M1-M2){n times}-M1 repeated patterns, suggesting that M1 and M2 mostly move together. However, the fact that M1 can be seen without M2, and is also more frequent (Table S3), suggests that they were once independent. It is not uncommon for MITEs to transpose alongside another MITE, like in the rice (*Oryza sativa*) genome where this event occurred several times (Tarchini et al. 2000) and where 11% of MITEs exist in multimers (Jiang and Wessler 2001). If M1 and M2 are genuine MITEs, they have to rely on an autonomous transposon for transposition. In *Pandoraviridae*, the *submariner* MITEs that colonized their genomes are related to a transposon present in their *A. castellanii* host genome (Sun et al. 2015; Zhang et al. 2018). We did not find transposons related to M1 and M2 in the available *A. castellanii* genome sequences, and current pithoviruses annotations show no sign of transposase. However, a recent study has found a putative transposase in the *Pithovirus sibericum* and *Pithovirus massiliensis* genomes, thanks to profile-based remote homology searches (Queiroz et al. 2023). The transposase found in *Pithovirus sibericum* is present within a repeat-rich region, while in *Pithovirus massiliensis*, it is adjacent to one. Those genes are thus candidates for explaining repeat invasion, but the fact that they are core genes, shared with all cedratviruses, makes it less likely. Here, we used structure-based remote homology searches thanks to Alphafold models coupled with Foldseek searches to further improve pithoviruses functional annotations. In doing so, we found candidate transposases that are systematically present in repeat-rich regions, and potentially undergoing pseudogenization (Table S5).

M1 and M2 repeats also share the characteristics of satellite DNA repeats, with a variable number of repeated units between closely related species, and a similar size range (Thakur et al. 2021). Furthermore, the pattern of higher sequence identity of repeats within the same regions (Fig. 5E) is reminiscent of the homogenization of repeats copies through concerted evolution (Thakur et al. 2021). This is particularly true for M1, but less so for M2 (Fig. S12). Thus, pithoviruses repeats might have also multiplied within genomes through recombination events rather than transposition, like in mice where it has been shown to increase the number of palindromic sequences (Zhou et al. 2001). In line with this hypothesis, our structure-based remote homology searches have revealed two proteins that could be involved in this process. The first one, pv_445 in *Pithovirus sibericum*, is probably a RuvC-like crossover junction endodeoxyribonuclease (Table S6). These proteins are involved in Holliday Junctions resolution formed by concatemers in *Nucleocytoviricota* with linear genomes such as *Poxviridae* (Culyba et al. 2009). They are also involved in the cleavage of cruciform (four-way junction) formed by inverted repeats, such as M1 and M2, to serve as intermediates in homologous recombination (Bowater et al. 2022). The second candidate, pv_159 in *Pithovirus sibericum*, is a probable SbcCD subunit D nuclease (Table S6). These proteins, in addition to the DNA Topoisomerase II (Froelich-Ammon et al. 1994) also found in *Pithoviridae*, are involved in the structural maintenance of chromosomes that cleave DNA hairpins and lead to homologous recombination (Connelly et al. 1998). The DNA breaks formed by these enzymes might also lead to the translocation of pithoviruses repeats into repeat-free regions.

It has been proposed that transposable elements can behave as seeds for the formation of satellite DNA (Meštrović et al. 2015; Garrido-Ramos 2017). The following scenario can thus be hypothesized for the invasion of pithoviruses’ genomes by repeats: initial transposition that occurred in the pithoviruses ancestor in one or several locations, probably followed by the inactivation of the transposase, and expansion by recombination as well as translocation into regions without repeats. The primo-invasion followed by drastic expansion occurred after the *Pithovirus*/*Cedratvirus* divergence. Was this the result of an explosive event or that of a gradual invasion remains to be determined. More deeply branching pithoviruses would be needed to settle this question. Interestingly, comparison of the *Pithovirus sibericum* and *Pithovirus massiliensis* genomes shows that the excision/insertion of their repeats has been ongoing since the two species diverged, more than 30,000 years ago (Legendre et al. 2014; Levasseur et al. 2016).

If repeat-rich regions constitute a large fraction of pithoviruses genomes, covering as much as a quarter of their genomes, they are also the source of genetic innovations. By comparing genes localized in repeat-rich regions with those in other regions, we found that they are more divergent and less evolutionary constrained (Fig. 6). We also found that repeat-rich regions are prone to gene capture of cellular and viral origins, and undergo many genomic rearrangements. One could hypothesize that the high conservation of repeat sequences triggers genomic recombination and gene exchange between co-infecting viral strains. As previously stated, our comparative analysis of the *Pithoviridae* family shows that they are conservative compared to other giant viruses’ families. These genomic islands might thus provide an opportunity for them to promote genetic diversity and raw genetic material for evolution to work on.

## Materials & Methods

### Isolation of cedratviruses

Cedratvirus borely and cedratvirus plubellavi were isolated in February 2017 from muddy soil samples from Marseilles, France (Parc Borély). The isolation and cloning of viruses were performed as previously described (Alempic et al. 2023). Briefly, mud was collected in sterile 50 mL Falcon tubes and several grams of this soil was resuspended, centrifuged and deposited on a Petri dish with 5000 *A. castellanii* (Douglas) Neff (ATCC 30010TM) cells/cm² growing on PPYG medium (for peptone, yeast extract and glucose). On the contrary of (Alempic et al. 2023), this step did not require fungizone. After three days, signs of a giant virus infections were looked for under the light microcope and amoeba cells were transferred to T25 cell culture flasks with *A. castellanii* cells in PPYG medium with ampicillin, cloramphenicol and kanamycin antibiotics and fungizone. After two to three days, when an ongoing viral infection was visible, cloning was performed by infecting a 6-well plate with 100,000 cells/cm² with an MOI of 2. After 1h30, the wells were washed 50 times with 2 mL of PPYG. Cells were then scraped and transferred to a 12-wells plate with 1 mL of PPYG in each well. Serial dilutions by 1/2 were performed. 0.4 µL from the 6th to 8th well were transferred to a 24-wells plate with PPYG medium. Wells with only one cell, as observed under the light microscope, were added with 200 cells then with 100,000 cells after three days. One week later, viruses were produced in T25 flasks with *A. castellanii* and purified on a cesium chloride gradient.

### Genome sequencing and assembly

250 ng of DNA of *Pithovirus mammoth* was sequenced by Oxford Nanopore, flowcell version R9.4.1 with the 1D native DNA barcoding protocol and by Illumina MiSeq. Long reads were basecalled by guppy v 2.1.3. Sequence data was assembled using a combination of short reads and long reads over 40kb by Unicycler v. 0.4.8 (Wick et al. 2017). The *Cedratvirus borely* genome was sequenced using Illumina MiSeq, assembled using Spades (version 3.9.1 and the “careful” option), scaffolded using SSPACE (SSPACE-LongRead v1.1) (Boetzer and Pirovano 2014) with Oxford Nanopore long reads (same protocol as above) and polished with Illumina reads using pilon v. 1.24. The *Cedratvirus plubellavi* genome was assembled using SPades (v 3.9.0) with Illumina MiSeq reads and Oxford Nanopore reads (option “careful”). Finally, the genomes of *Cedratvirus lena* and *Cedratvirus duvanny* were sequenced using Illumina MiSeq and NovaSeq technologies. *Cedratvirus lena* was assembled after removing reads mapped to a contaminant *Pandoravirus* using Bowtie 2. *Cedratvirus lena* and *Cedratvirus duvanny* reads were trimmed by BBduk (<u>sourceforge.net/projects/bbmap/</u>) and assembled using SPAdes v 3.14 (Prjibelski et al. 2020) with options “careful” and k=15,17,19,21,29,33,41,55,63,71,91,101,115. The scaffolding was then performed by RaGOO (Alonge et al. 2019) using *Cedratvirus kamchatka* as template. The genome of *Pithovirus sibericum* was reassembled using PacBio long reads over 500 bp from (Jeudy et al. 2020) with Unicycler v 0.5.0. 100000 sampled short reads pairs from (Legendre et al., 2014) were trimmed with BBduk and aligned to the genome with bowtie2 with option “no-discordant”. Polishing was done by pilon v. 1.24.

The 10 metagenomic datasets with highest density of reads matching the M1 or M2 repeats (Table S4), were retrieved from the SRA archive and cleaned using BBduk with the following parameters: “ref=adapters ktrim=r k=23 mink=11 hdist=1 tpe tbo”. The datasets were then individually assembled using megahit (v 1.1.3) with default parameters.

The 3 pithoviruses and the 9 cedratviruses genomic sequences (accessions in Table S2A and S2B) were then artificially linearized to start at the same position for comparative analyses. *Cedratvirus A11* and *Pithovirus sibericum* were used as reference to linearize cedratviruses and pithoviruses. All genomes were aligned with progressiveMauve and visualized with Mauve to identify the corresponding starting positions in other *Pithoviridae*. Their genomes were then cut and swapped at this reference position. The genome of *Brazilian cedratvirus,* having undergone several genomic rearrangements, was then reverse complemented to better fit the genome structure of other cedratviruses (Fig. S4).

### Genome annotation

For functional annotation, genes were predicted using Genemark (Besemer et al. 2001) with option – virus. ORFs over 50 amino acids were kept for publication and ORFs over 100 amino acids were used for core- and pan-genome comparative analyses.

ORFs were annotated using InterProScan (v5.39-77.0, databases PANTHER-14.1, Pfam-32.0,ProDom-2006.1, ProSitePatterns-2019_01, ProSiteProfiles-2019_01, SMART-7.1, TIGRFAM-15.0) (Jones et al. 2014) and CDsearch (Conserved Domain Database) (Lu et al. 2020) with default options. We also searched for viral specific functions using hmmsearch on the virus orthologous groups database (https://vogdb.org/) from November 2022 with an *E*value cutoff of 10^-5^. ORFs were compared to the nr and swissprot databases using BLASTP (Altschul et al. 1990) and *E*values cutoff of 10^-2^. Transmembrane domains were identified with Phobius (Käll et al. 2004).

For further improving the functional annotation, we generated Alphafold2 (Jumper et al. 2021) models of the proteins of all three pithoviruses using the Colabfold (Mirdita et al. 2022) pipeline version 1.52. Multiple Sequence Alignments were computed using MMSeqs2 (Steinegger and Söding 2017) with default parameters and the UniRef30 (version 2202), pdb70 (version 220313) and envDB (version 202108) databases. Structure prediction was processed with default parameters, automatic number of recycles (up to 20), no templates, and no relax. For each protein, only the top ranked model (by pTM) was selected for the next steps. We next used Foldseek (van Kempen et al. 2023) through API to search the alphafold-swissprot, alphafold-proteome and PDB100 databases from July 2023. For final annotations in *Pithovirus sibericum* (Table S6), only matches with over 50% of subject coverage, an E-value below 0.01 and a probability over 50% were considered. The consistency in-between the function of the best Foldseek matches was checked and functional annotation was done manually.

Two-way AAI were calculated for all pairs of genomes by the package enveomics (Rodriguez-R and Konstantinidis 2016) and options --id 15 -L 0.4.

Relative synonymous codon usage was calculated using an in-house script, genome_metrics.py, for fast genome-wide analysis relying on Biopython (Cock et al. 2009). Given one amino-acid *a* and a codon *c*, we applied the following formula:

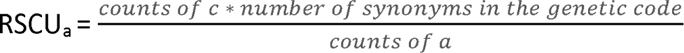

The heatmap and multi-dimensional scaling were done on the RSCU calculated on the whole genome (i.e treating all the codons in the genome at once) excluding stop codons and tryptophan codons.

### Computation of orthologous gene groups and phylogeny

A phylogenetic tree was computed by OrthoFinder (v2.5.4) (Emms and Kelly 2019) using all available *Pithoviridae* genomes in addition to the *Orpheovirus* (Andreani et al. 2018), *Hydrivirus* (Rigou et al. 2022) and *Marseillevirus* genomes (Table S2). The tree was then rooted using the distantly related *Marseillevirus* (Boyer et al. 2009) as an outgroup. Hierarchical Orthologous Groups (HOGs) were then determined by OrthoFinder (v2.5.4) using this rooted tree. A final phylogeny was inferred on the concatenated alignment of single copy core HOGs by IQ-TREE (Nguyen et al. 2015) with the LG+F+G4 model and options -bb 5000 -bi 200.

The distribution of core genes and duplications along the *Pithoviridae* genomes were evaluated through a sliding window of 21 genes. The genomes were considered circular; meaning that, at the extremities of the artificially linearized genomes, the window would span across the other end of the genome.

Selection pressure on genes was estimated by the ratios of non-synonymous substitution rates (dN) to synonymous substitution rate (dS), calculated by codeml of the PAML v4.9 package (Yang 1997). All pairs of single copy orthologues as defined by OrthoFinder were retrieved and aligned with T-Coffee (Notredame et al. 2000). Codeml was given the sequence pairs alignments and the resulting dN/dS ratio was considered only if dS < 1.5, dS > 0.1 and dN/dS < 10. Later, the dN and dS values for each gene was estimated as the mean of all value calculated on gene pairs.

### Estimation of the core and pan-genomes of cedratviruses

Core/pan-genomes sizes were calculated on HOGs (Hierarchical Orthologous Groups) at the root node. Genomes were iteratively added with all possible combinations to simulate a dataset with 1 to 9 genomes. We used the presence/absence matrix of HOGs instead of gene counts as in the original method (Tettelin et al. 2005). Data were processed using R (v4.04 (R Core Team 2021)). First, OrthoFinder data was transformed into a numeric matrix with the function “HOG2df” taking as arguments Phylogenetic_Hierarchical_Orthogroups/Nx.tsv and Orthogroups_UnassignedGenes.tsv file locations. Next, the simulated datasets were processed by functions “get_core_pan_new_info_Real” for cedratviruses (Fig. 3A) and with function “get_core_pan_info” for the comparative analyses with normalized genome sizes (Fig. 3B and C). In addition to this method, the micropan package (Snipen and Liland 2015) was used to estimate the closedness/openness of the different pan-genomes applying heap’s law with option n.perm = 1000. Pan-genomes are considered open when the estimated α parameter < 1 and closed otherwise.

For comparison, the ORF predictions, orthology analyses and core/pan-genome estimations were performed on other viral families: *Pandoraviridae* (Table S2C), *Marseilleviridae* (Table S2D), ranaviruses (Table S2E), *Megavirinae* (Table S2F). The outgroups used were respectively *Mollivirus sibericum*, *Ambystoma tigrinum virus*, *Red seabream iridovirus* and *Chrysochromulina ericina virus*.

### Identification of horizontal gene transfers

HGTs were identified based on phylogenetic trees of each HOG complemented with homologous sequences that were retrieved using a two steps procedure. First, the sequences of each HOG were aligned using DIAMOND BLASTP (Buchfink et al. 2015) against the RefSeq database from March 2019 (O’Leary et al. 2016) with an *E*value threshold of 10^-5^, keeping only matches covering more than 50 % of the query. Up to 10 matches per domain (Bacteria, Archaea, Eukaryota and Viruses) were kept for each query and CD-hit was applied on the retrieved sequences. Secondly, the resulting sequences were queried again against the RefSeq using DIAMOND with the same *E*value threshold. A maximum of two proteins per domain, whose matches covered more than 80 % of the query, were kept at this point. The HOGs and selected sequences from the first and second rounds were aligned using MAFFT v7.475 (Katoh and Standley 2013) and phylogenetic trees were built using IQ-TREE with options -bb 1000 -bi 200 -m TEST. Each resulting phylogenetic tree was rooted by mad v2.2 (Tria et al. 2017). Trees were finally visually inspected and HGT events counted when one or several *Pithoviridae* genes were within a bacterial, eukaryotic, archaeal or different viral clade.

### Detection and classification of genomic repeats

A pipeline was developed to retrieve repeat-rich regions and map individual repeats from pithoviruses’ genomes. The steps were: (1) genome-wide alignment, (2) flattened dotplot calculation, (3) repeat-rich regions mapping, (4) individual repeats retrieval, (5) repeat clustering.

1. genomes were aligned against themselves by BLASTN with an *E*value threshold of 10^-10^.
2. for each position of the genome, the number of times it was aligned was counted resulting in a vector (y); similar to a flattened dotplot.
3. A smooth vector (y_Ss_) was first estimated by sliding mean filtering with a window size of 500 nt. A detection threshold (τ) was calculated as 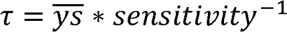, with a sensitivity coefficient set to 2.5. Repeat-rich regions were detected by comparing the vector y_s_ to τ. Repeat-rich regions were defined as regions where y_s_ is above the threshold τ. Each region’s start and stop are thus the positions of intersections of y_s_ and τ.
4. For each previously detected region, individual repeats were extracted using a smoothed derivative of y. Smoothing was applied before and after the derivation, this time with a window size of 20 nt. Then, the absolute value was taken in order to obtain the vector |y_s_|. Then the local maxima were considered as repeat delimitations if above a cutoff set to 10.
5. repeats are globally aligned to each other by needle of the EMBOSS suite (Rice et al. 2000). They are then ordered according to the mean distance (100 – needle identity percentage) to their 10 closest neighbors. The first sequence becomes a reference sequence. Then, sequences are clustered together if they are at least 70 % identical to a reference or they become themselves a reference. Finally, clusters are merged together if over half of their respective sequences are at least 70 % identical. For visual inspection to infer repeat types and similarity in-between clusters, a matrix of dotplots presenting the alignments of reference sequences is drawn.

For an in-depth analysis of pithoviruses’ repeats, the sequences from the largest cluster of repeats (M1) were aligned with MAFFT and trimmed according to the position of the aligned terminal “TA”. The reference sequence (see step 5) of M1 and M2 were folded by mFold (Zuker 2003). To retrieve divergent M1 and M2 clusters, the dotpots of reference sequences was visually inspected. Reference sequences aligned to the reference of M1 or M2 clusters were annotated as M1 or M2-like (example given by cluster 3 in step 5, Fig. S7).

MUST v2-4-002 (Ge et al. 2017) and MITE-Tracker (Crescente et al. 2018) were used to infer the nature of the repeats.

For comparison we also ran RepeatModeler v2.0.4 on the genome of *Pithovirus sibericum*.

### Repeat and adjacent sequences similarity

To compare the similarity of M1 or M2 within the same or different regions, we used the percentage of identity calculated from pairwise global alignments by needle in step 5 of the described pipeline. To compare similar numbers of pairs, we randomly sub-sampled pairs of repeats originating from different repeat-rich region to match the number of pairs of repeats originating from the same region. The operation was done several times and the results were always comparable to the results in Fig. 5.

### Statistics of genes within repeat-rich regions

The repeat-detection pipeline was used with a smoothing window size of 4000 nt (3) to define repeat-rich regions. Bedtools intersect was used to reveal the genes that were within repeat-rich regions. GO term enrichment analyses were performed on those genes by extracting the GO terms of the non-overlapping protein domains predicted by InterProScan in the three genomes. The topGO package of R was used to test the significantly enriched molecular functions in repeat-rich regions versus outside those regions through a Fisher test.

## Acknowledgments

This work was supported by the Agence Nationale de la Recherche grant (ANR-22-CE12-0041) to ML, (ANR-10-INBS-09-08) to J-MC and CNRS Projet de Recherche Conjoint (PRC) grant (PRC1484-2018) to CA. S.R. was supported by a doctoral fellowship obtained from Aix-Marseille University. We thank the PACA Bioinfo platform for computing support and the GENCI-IDRIS for the GPU HPC resources used for Alphafold predictions (Grant 2022-AD011013526). We also thank Hugo Bisio for carefully reading the manuscript, as well as Bernard La Scola and Julien Andreani for giving access to the raw sequencing data of Brazilian cedratvirus and Victória Queiroz for providing additional gene annotations.

## Availability of Data and Materials

Genome sequences and annotations of the following five *Pithoviridae* have been deposited to GenBank: *Cedratvirus borely* (OQ413575), *Cedratvirus plubellavi* (OQ413576), *Cedratvirus lena* (OQ413577, OQ413578, OQ413579, OQ413580), *Cedratvirus duvanny* (OQ413581) and *Pithovirus mammoth* (OQ413582).

R functions for pan- and core-genome analysis as well as HOGs are available for download: https://doi.org/10.6084/m9.figshare.23913051 together with rooted trees and the final HGT analysis results. Additional in-house scripts are also provided here.

The code for pithovirus repeats detection and clustering is available at: https://src.koda.cnrs.fr/igs/genome-repeats-detection.git.

None of the authors have any competing interests.

