## Supplementary material for "Pithoviruses are invaded by repeats that contribute to their evolution and divergence from cedratviruses"

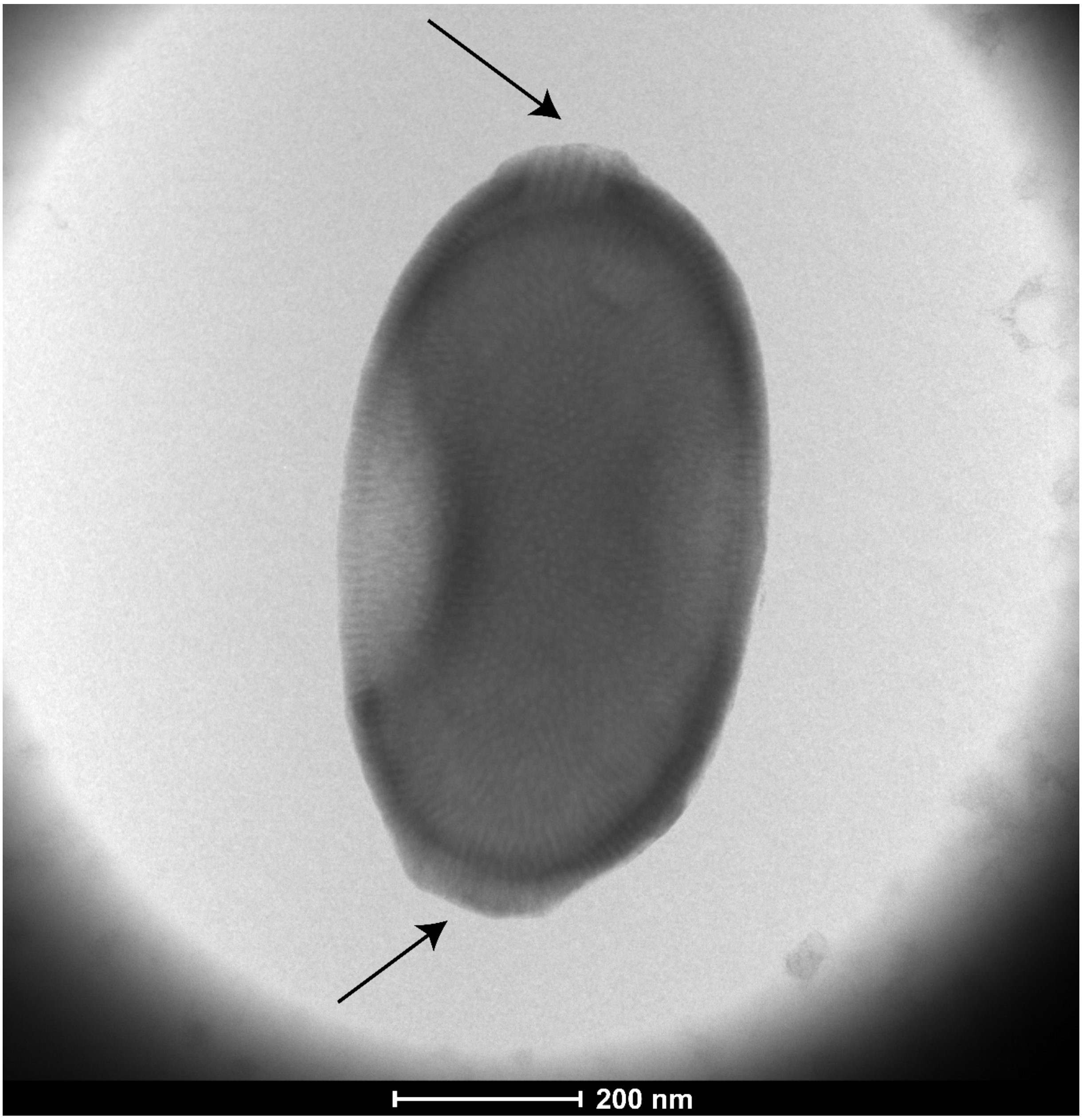


**Figure S1. Negative staining microscopy of cedratvirus plubellavi**
Corks at each apex of the viral particle are shown with arrows.

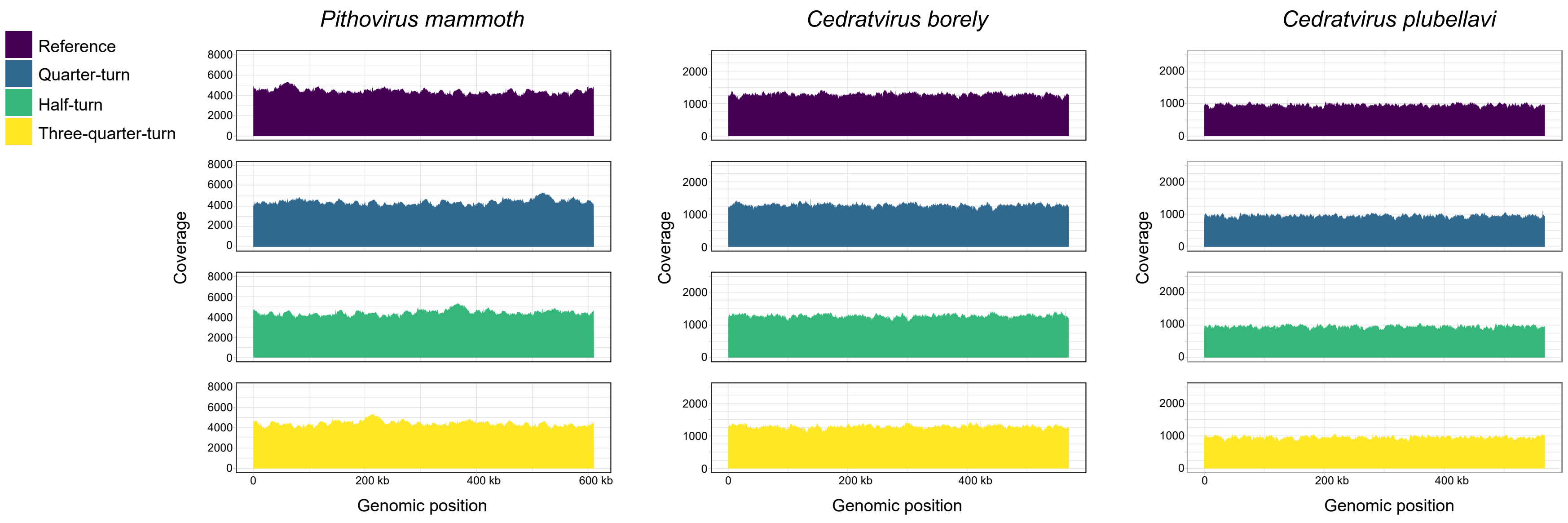


**Figure S2. Long reads coverage along *Pithoviridae* genomes linearized at 4 different positions**

The assembled genome of *Pithovirus mammoth* (left), *Cedratvirus borely* (center) and *Cedratvirus plubellavi* (right) were linearized at four equidistant positions and reads were mapped on these references. The ONT read coverage along these genomes is shown.

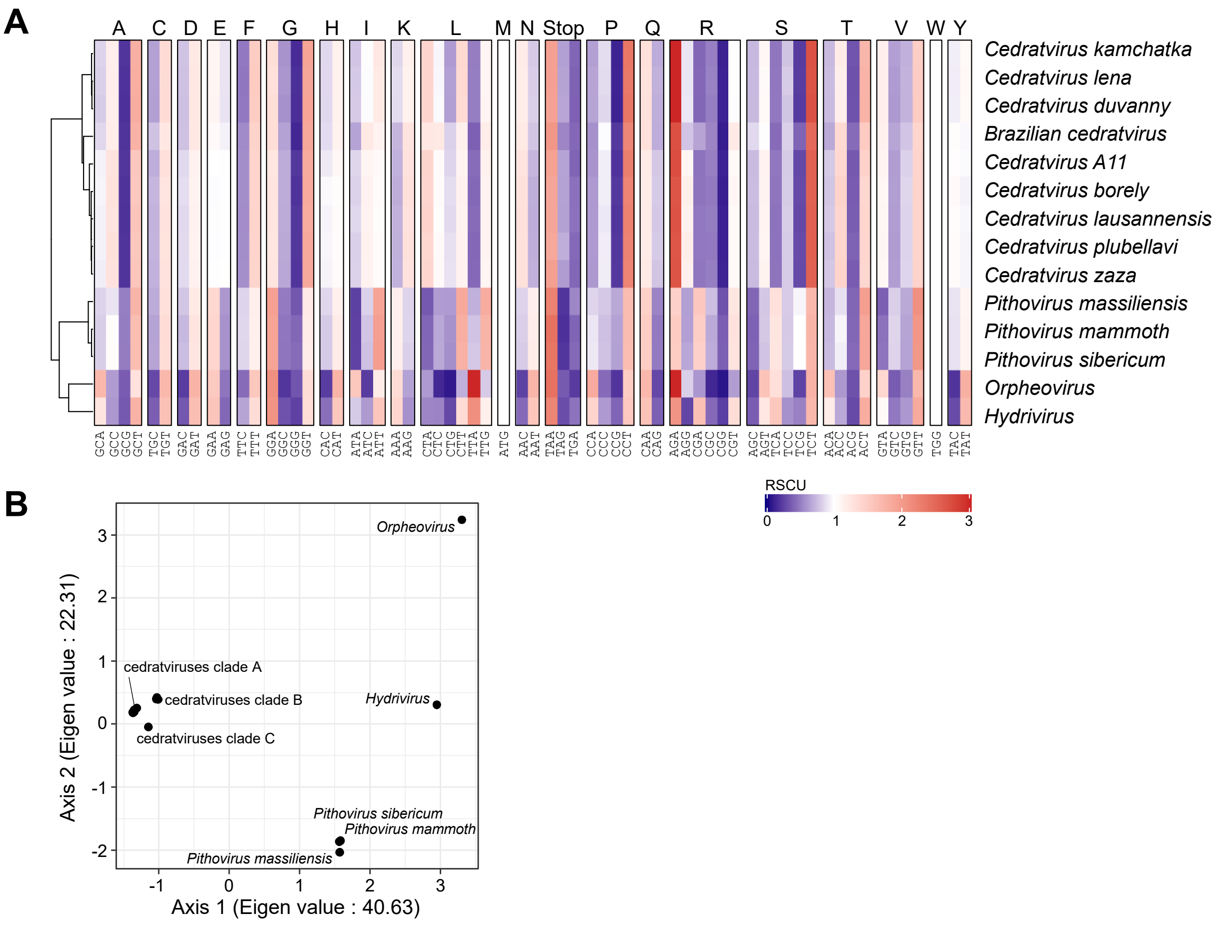


### Figure S3. Comparison of relative synonymous codon usages

(A) Codon usage bias for each amino acid represented by the RSCU value. (B) PCOA analysis of the RSCU values without the stop codons, the tryptophan and the methionine codons.

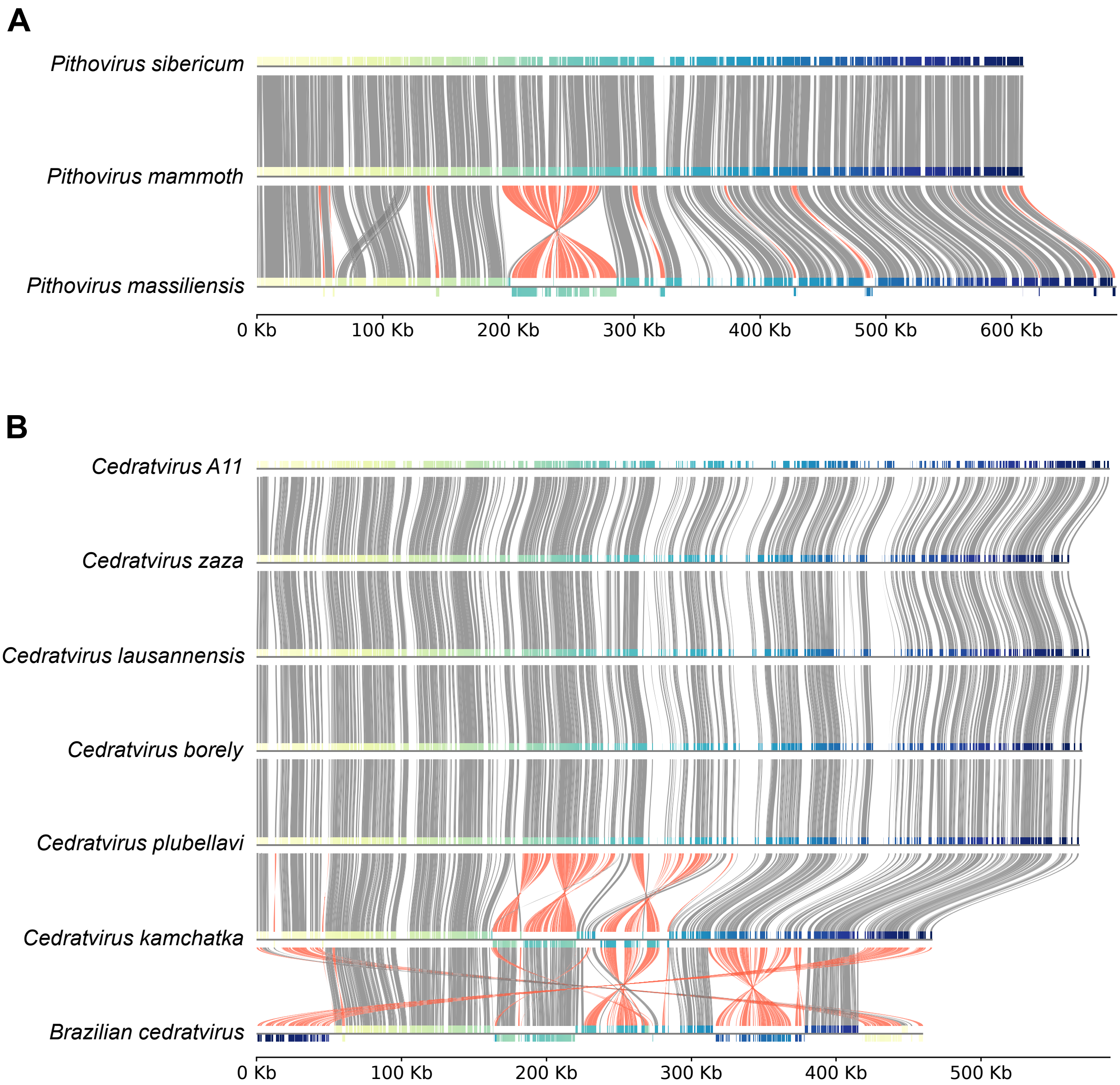


**Figure S4. Genome alignment of *Pithoviridae***

Shared nucleotide sequence blocks within clades were drawn based on the alignment by progressive-mauve (Darling et al. 2010) of (A) the three pithoviruses and (B) seven cedratviruses. *Cedratvirus lena* and *Cedratvirus duvanny* have been excluded since the assembly was incomplete (multiple contigs). Syntenic regions are shown in gray and large inversions in red. ORFs are color-coded from yellow to blue according to their genomic positions. A scale bar of genome sizes is shown at the bottom.

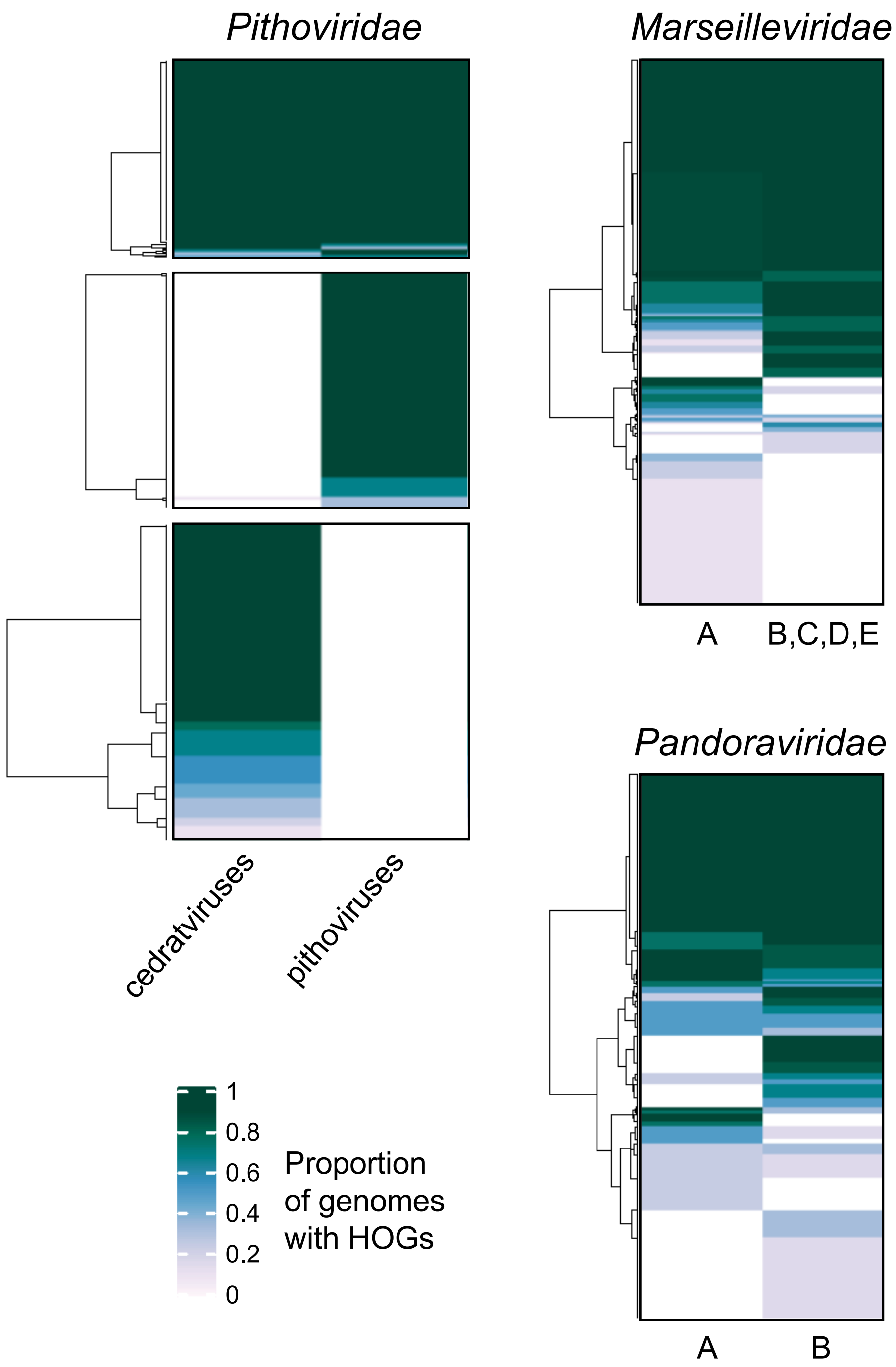


**Figure S5. Patterns of presence/absence of HOGs within viral family's sub-clades**
Species were divided into clades ignoring the outgroup. The number of species from each clade that appeared in each HOG was then counted.

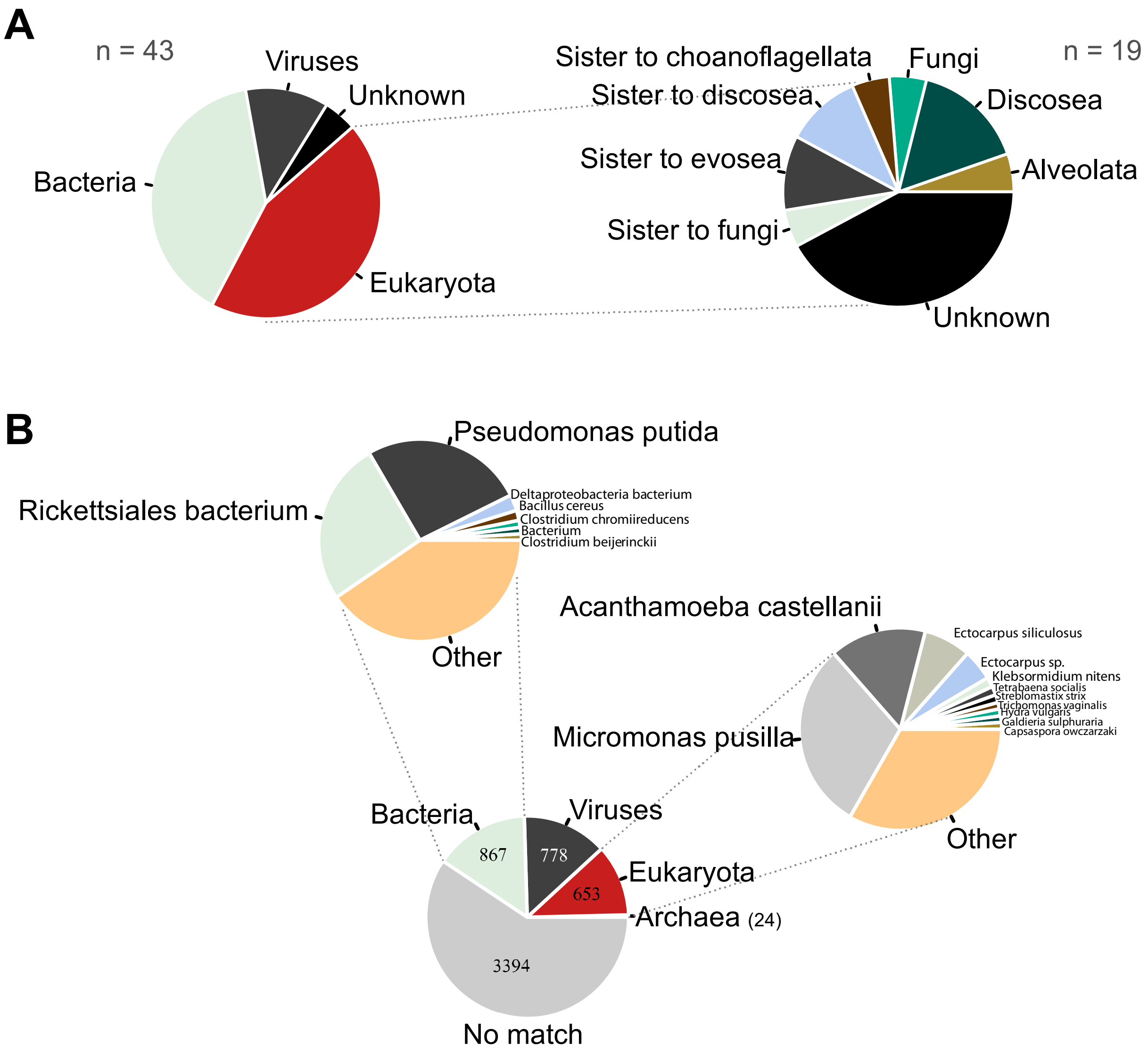


**Figure S6. Horizontal Gene Transfer events in *Pithoviridae* and BLASTP control**
For each HGT event, the likely origin as estimated from the visualization of phylogenetic trees (A) and best BLASTP results (*E*value ≤ $10^{-5}$) from the nr database free of *Pithoviridae* (B) are shown. From eukaryotes, “sister to” is short for “sister group of…”. Bacterial and eukaryotic species with more than 1% of matches in their respective category are shown.

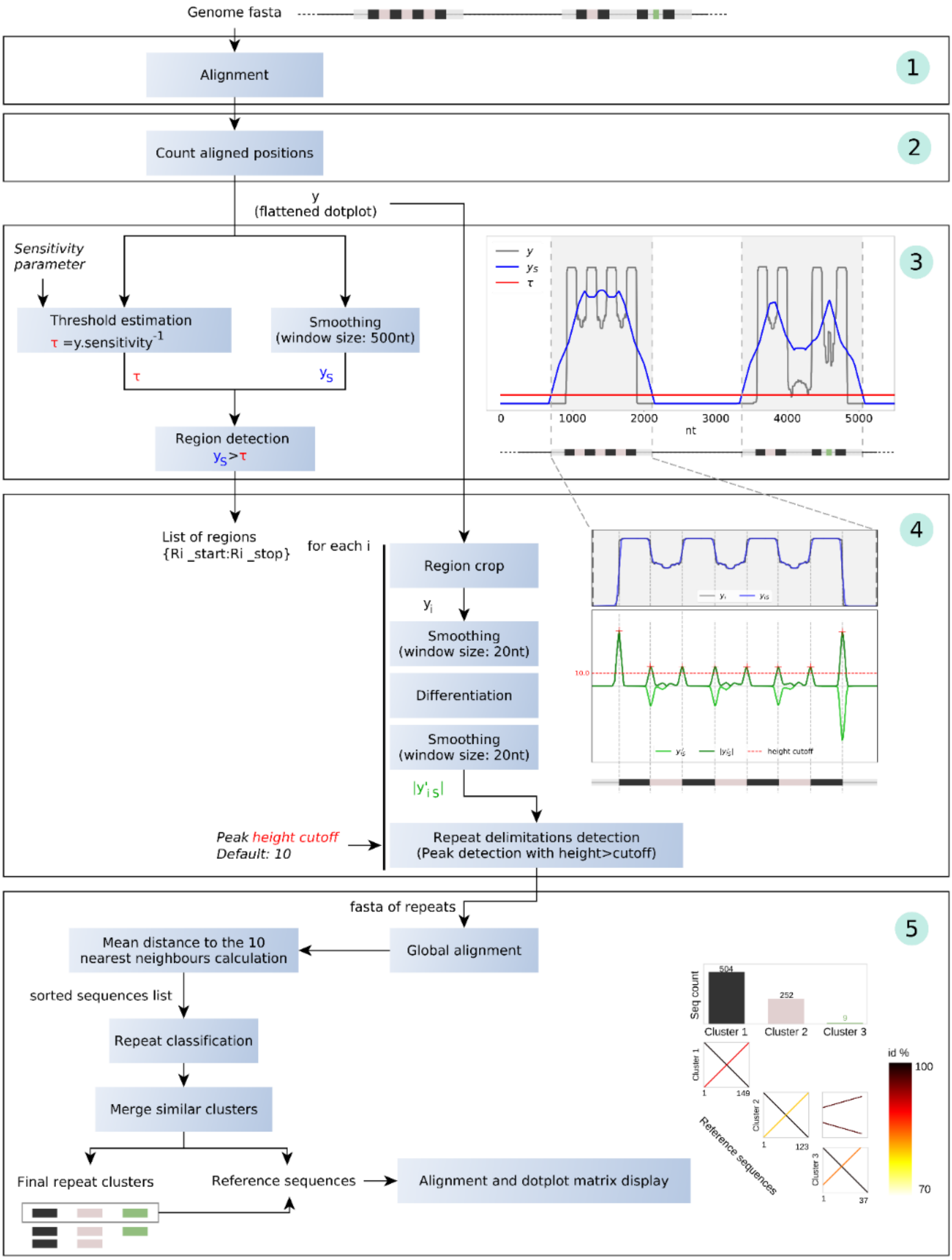


**Figure S7. Workflow for repeat analysis**

Steps one to five are represented within large boxes. Operations are in blue boxes while objects are shown as black text. Besides "Genome fasta" is schematized a portion of the genome containing repeats as colored boxes. The slightly grey boxed represent unclustered sequences.

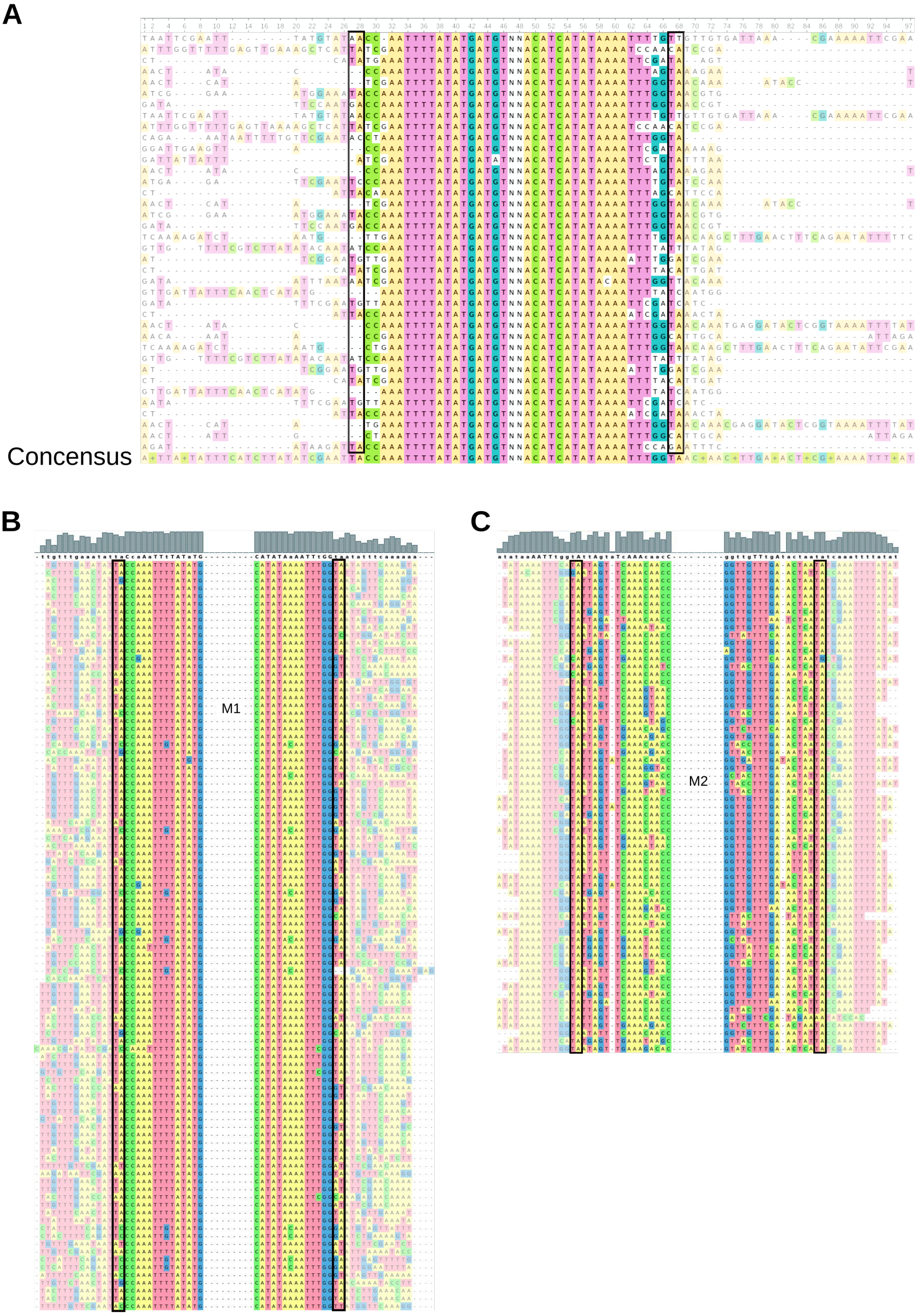


**Figure S8. Alignment of repeats and adjacent sequences**

(A) 40 bp around the beginning and end of repeat-rich regions in *Pithovirus sibericum* were aligned with MAFFT and visualized with UGENE. Upstream and downstream regions are in shaded colors while repeat regions are in plain colors. The center of each repeat region is truncated and indicated with "NN". Boxes indicate putative TSD. (B) M1 and (C) M2 reference sequences without the surrounding TA were aligned to the genome of *Pithovirus sibericum.* The matching sequences, extended by 15 bp were aligned. The mid-part of the alignments were truncated and indicated with "-". Black boxes indicate the proposed TSD sites.

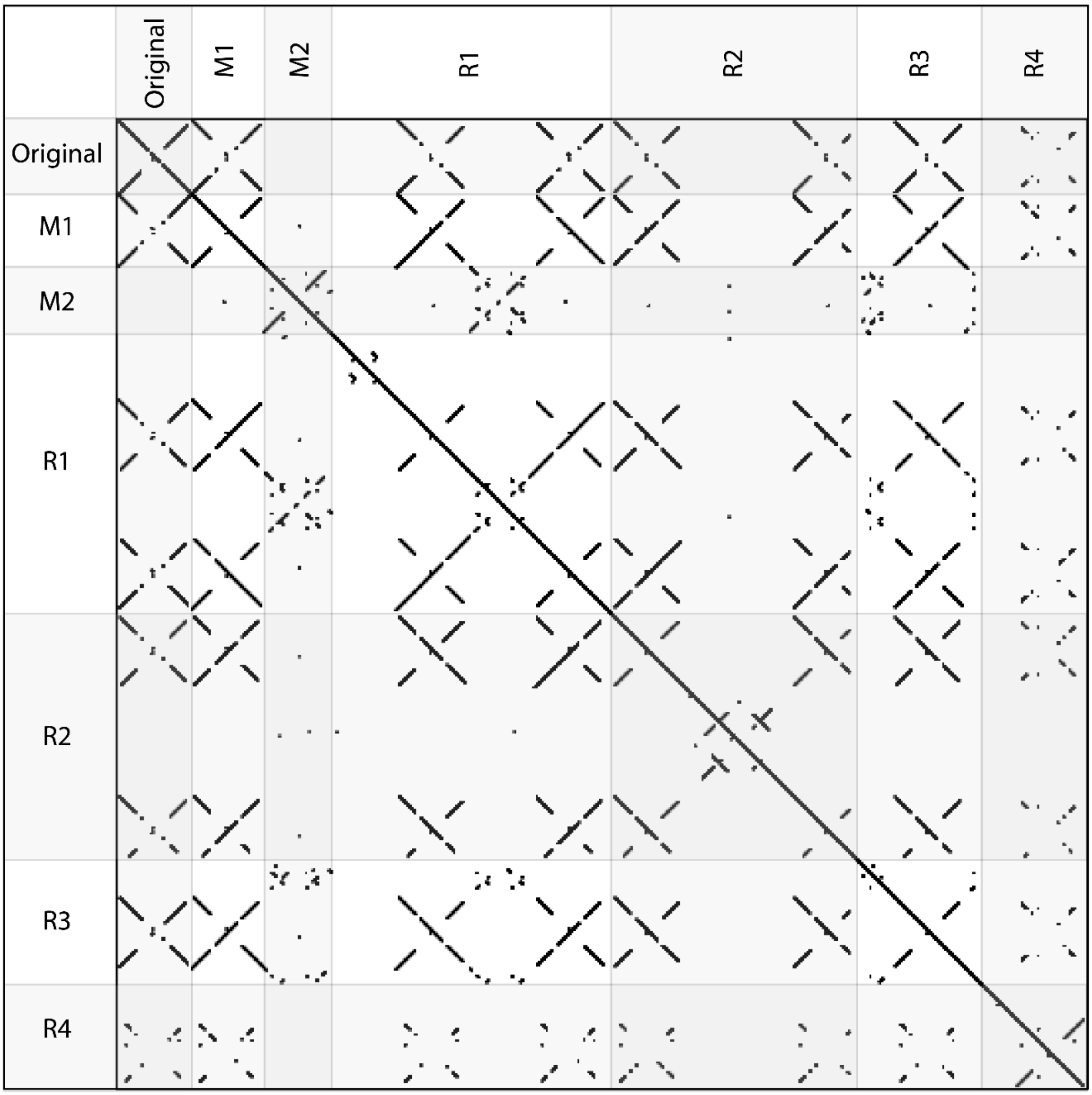


**Figure S9. Dotplot of consensus repeats sequences found in pithoviruses**

The dotplot includes the repeat previously identified in *Pithovirus sibericum* (Legendre et al. 2014) coined "Original", as well repeats identified by our dedicated pipeline (M1 and M2) and the ones identified by RepeatModeler version 2.0.4 (R1, R2, R3 and R4).

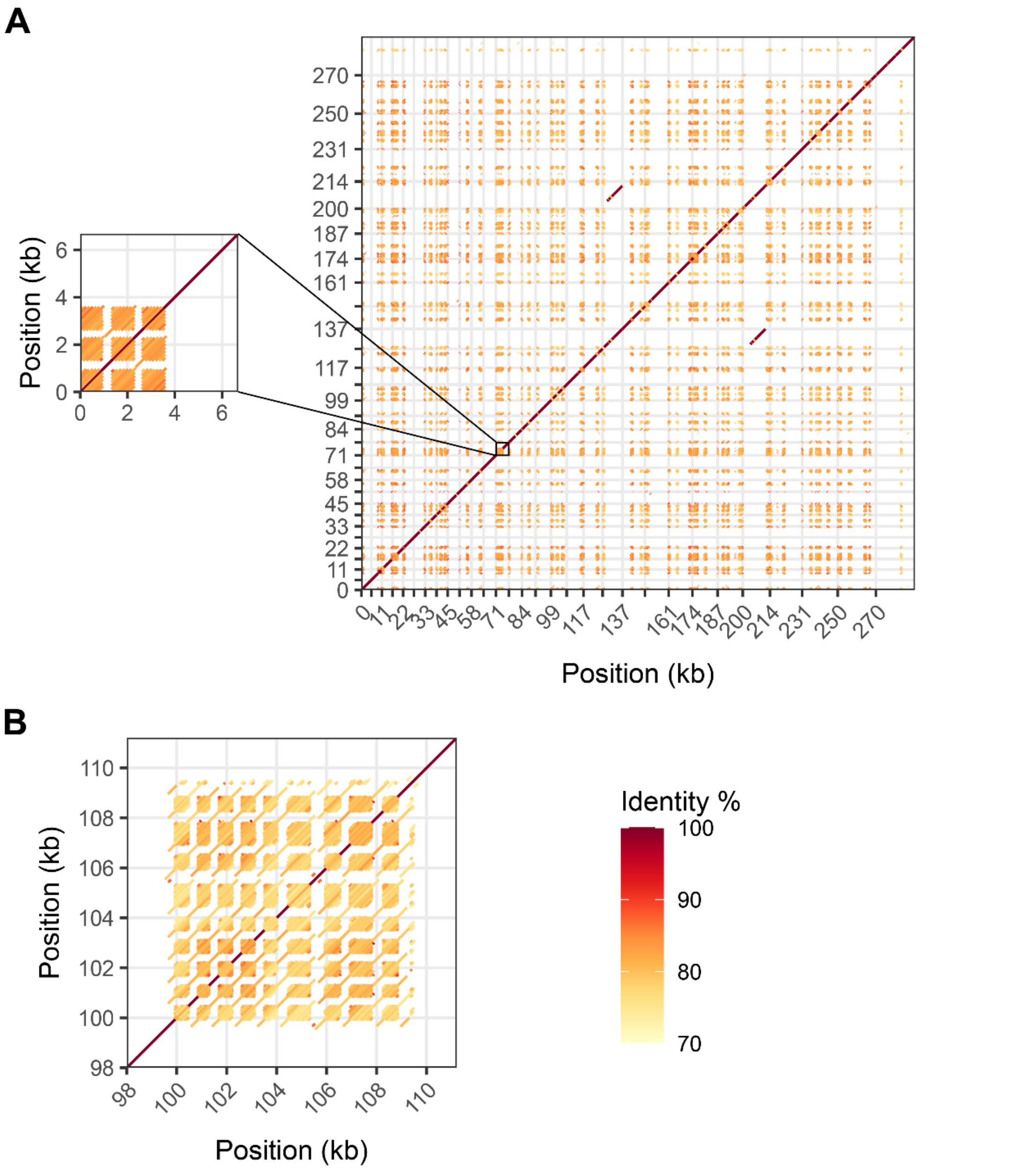


**Figure S10. Repeats found in *Pithoviridae*-like metagenomes**
(A) *Pithovirus LCPAC302* (Bäckström et al. 2019) presents numerous direct repeats. In some rare cases, these repeats are interspersed by a similar sequence as shown in the inset. X-axis and y-axis breaks correspond to the delimitation of contigs. (B) Regularly interspersed direct repeats from a permafrost *Pithoviridae*-like metagenome (K_bin2137_k1) (Rigou et al. 2022).

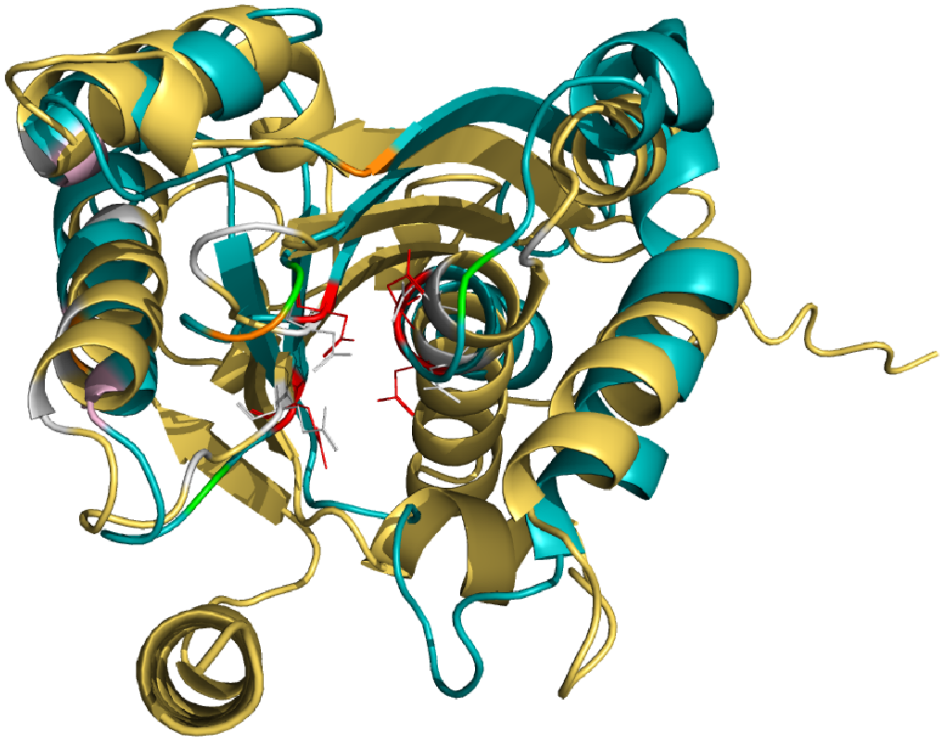

| Fowlpox virus (PDB: 6P7B) | Pithovirus pv_445 |
|---|---|
| DDE site | |
| D7 | D57 |
| E60 | E115 |
| D132 | D200 |
| N/D135 | D203 |
| DNA cleavage | |
| N12 | K62 |
| Q62 | Q117 |
| K129 | K197 |
| DNA binding | |
| K11 | K60 |
| K34 | |
| K70 | R126 |
| DNA contact | |
| W41 | F92 |
| P67 | P122 |
| K70 | R126 |
| Y73 | 7Y134 |

**Figure S11. Superposition of the *Fowlpox virus* Holliday Junction resolvase protein structure and the Alphafold model of the *Pithovirus sibericum* pv_445 homolog**

Shown on the left is the superposition of the *Fowlpox virus* protein structure (PDB 6P7B) in blue and the Alphafold structure model of *Pithovirus sibericum* pv_445 in yellow. Important residues from the *Fowlpox virus* structure (Li et al. 2020) are color-coded and described in the table on the right. Corresponding residues in the *Pithovirus sibericum* structure model are shown in gray and described in the table as well.

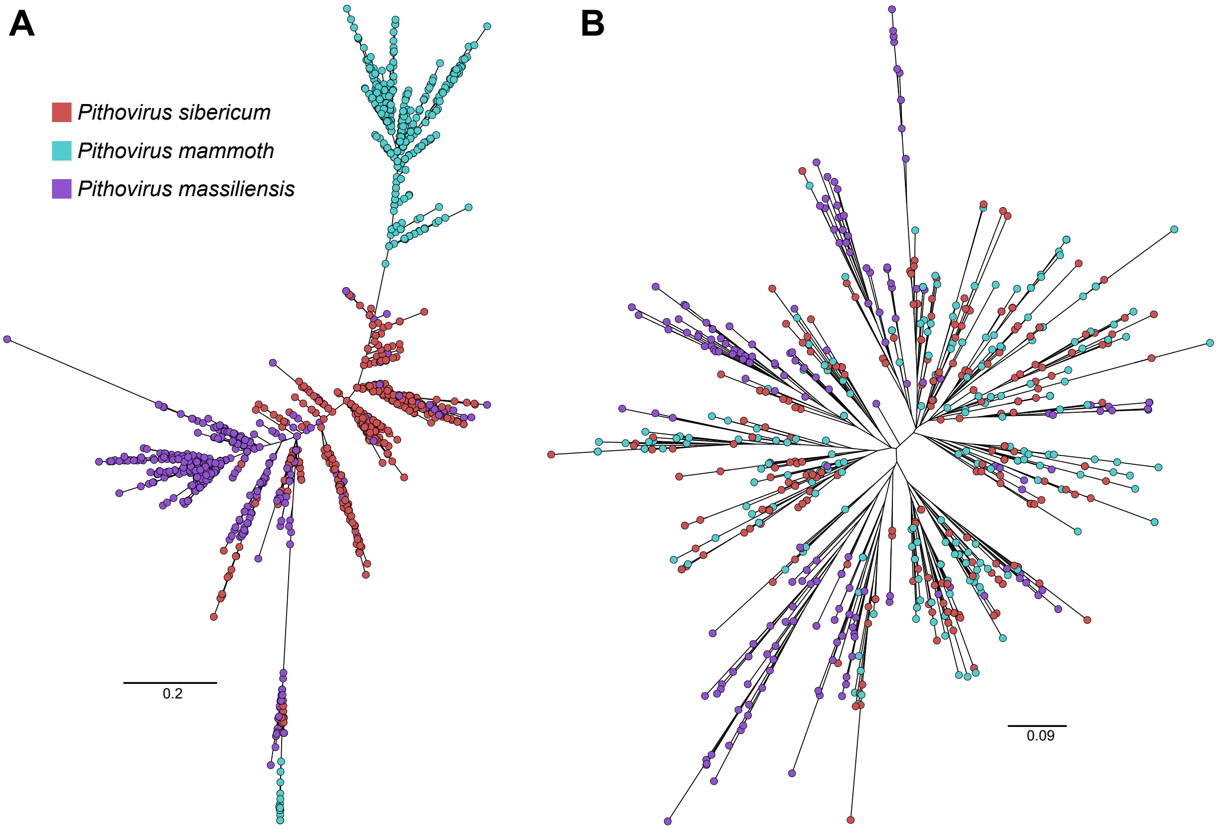


**Figure S12. Phylogeny of the M1 and M2 repeats of three pithoviruses**
Phylogenetic tree of the M1 (A) and M2 (B) units computed by IQtree with best-fit models TPM2+F+I+G4 and TVM+F+I+G4, respectively. The alignment of the units after correction of strand orientation was performed using Mafft with options "--maxiterate 1000 --localpair".

**Table S1. Statistics for the genome assemblies of *Pithoviridae***

The genomes were assembled with a combination of long and short reads, and with short reads only for comparison. For all assemblies we counted the length of all types of repeats altogether, not only M1 and M2.

| | *Pithovirus mammoth* | | *Cedratvirus borely* | | *Cedratvirus plubellavi* | |
|---|---|---|---|---|---|---|
| | Illumina | Illumina +ONT | Illumina | Illumina +ONT | Illumina | Illumina +ONT |
| # contigs > 1 kb | 42 | 2 | 3 | 1 | 1 | 1 |
| # contigs > 2 kb | 35 | 1 | 3 | 1 | 1 | 1 |
| # contigs > 5 kb | 27 | 1 | 3 | 1 | 1 | 1 |
| # contigs > 10 kb | 20 | 1 | 3 | 1 | 1 | 1 |
| # contigs > 25 kb | 7 | 1 | 3 | 1 | 1 | 1 |
| # contigs > 100 kb | 0 | 1 | 2 | 1 | 1 | 1 |
| Max contig length (kb) | 42 | 610 | 305 | 570 | 566 | 568 |
| Total repeats (bp) | 74224 | 150628 | 3056 | 15706 | 9825 | 11712 |
| Total repeats (%) | 14.1 | 24.7 | 0.5 | 2.8 | 1.7 | 2.1 |

### Table S2. Assemblies used for comparative genome size analysis

| **A) Previously published** | **NCBI accessions** | **E) Ranaviruses** | |
|---|---|---|---|
| *Pithovirus sibericum* | NC_023423.1 | *Ambystoma tigrinum virus* | GC_ 000841005.1 |
| *Pithovirus massiliensis* | SAMEA4074172 | *Bohle iridovirus* | GCF_002826565.1 |
| *Cedratvirus A11* | NC_032108.1 | *Common midwife toad virus* | GCF_003033105.1 |
| *Cedratvirus lausannensis* | LT907979.1 | *Epizootic haematopoietic necrosis virus* | GCF_000897115.1 |
| *Cedratvirus zaza* | LT994652.1 | *European catfish virus* | GCF_000897115.1 |
| *Brazilian cedratvirus* | LT994651.1 | *Frog virus 3* | GCF_001717415.1 |
| *Cedratvirus kamchatka* | MN873693.1 | *Infectious spleen and kidney necrosis virus* | GCF_000848865.1 |
| *Orpheovirus* (outgroup) | NC_036594.1 | *Lymphocystis disease virus 1* | GCF_000839605.1 |
| *Hydrivirus* (outgroup) | GCA_943296135.1 | *Lymphocystis disease virus-isolate China* | GCF_000844885.1 |
| *Marseillevirus* (outgroup) | NC_013756.1 | *Lymphocystis disease virus Sa* | GCF_001974475.1 |
| | | *Ranavirus maximus* | GCF_001717415.1 |
| **B) New *Pithoviridae*** | | *Largemouth bass virus* | GCA_013122655.1 |
| *Cedratvirus borely* | OQ413575 | *Scale drop disease virus* | GCF_001274405.1 |
| *Cedratvirus plubellavi* | OQ413576 | *Short-finned eel ranavirus* | GCF_001678255.2 |
| *Cedratvirus lena* | OQ413577<br>OQ413578<br>OQ413579<br>OQ413580 | *Singapore grouper iridovirus* | GCF_000846905.1 |
| *Cedratvirus duvanny* | OQ413581 | *Grouper iridovirus* | GCA_006465545.1 |
| *Pithovirus mammoth* | OQ413582 | *Red seabream iridovirus* (outgroup) | GCA_011894875.1 |

| **C) *Pandoraviridae*** | | **F) *Megavirinae*** | |
|---|---|---|---|
| *Pandoravirus braziliensis* | LT972217.1 | *Acanthamoeba polyphaga lentillevirus* | GCA_000320725.1 |
| *Pandoravirus celtis* | MK174290.1 | *Mamavirus* | GCA_002966335.1 |
| *Pandoravirus dulcis* | GCA_000911655.1 | *Megavirus chilensis* | GCF_000893915.1 |
| *Pandoravirus inopinatum* | GCA_000928575.1 | *Megavirus courdo7* | GCF_000893915.1 |
| *Pandoravirus macleodensis* | GCA_003233935.1 | *Megavirus vitis* | GCA_004156275.1 |
| *Pandoravirus massiliensis* | MZ384240.1 | *Mimivirus* | GCA_024266865.1 |
| *Pandoravirus neocaledonia* | GCA_003233915.1 | *Moumouvirus australiensis* | GCA_004156295.1 |
| *Pandoravirus pampulha* | OFAJ00000000.1 | *Moumouvirus* | GCF_000904035.1 |
| *Pandoravirus quercus* | GCA_003233895.1 | *Tupanvirus deep ocean* | GCA_002966475.2 |
| *Pandoravirus salinus* | GCA_000911955.1 | *Tupanvirus soda lake* | GCA_002966485.2 |
| *Mollivirus sibericum* (outgroup) | NC_027867.1 | *Chrysochromulina ericina virus* (outgroup) | GCF_001399245.1 |

| **D) *Marseilleviridae* as in** (Blanca et al. 2020) | |
|---|---|
| *Marseillevirus* | GU071086 |
| *Lausannevirus* | HQ113105 |
| *Cannes 8 virus* | KF261120 |
| *Insectomime virus* | HG428764 |
| *Tunisvirus* | KF483846 |
| *Brazilian marseillevirus* | KT752522 |
| *Melbournevirus* | KM275475 |
| *Port-miou virus* | KT428292 |
| *Tokyovirus* | Reassembled in (Blanca et al. 2020) |
| *Noumeavirus* | KX066233 |
| *Golden marseillevirus* | KT835053 |
| *Kurlavirus* | KY073338 |
| *Marseillevirus shanghai* | MG827395 |
| *Ambystoma tigrinum virus* (outgroup) | MK580533.2 |

**Table S3. Pithoviruses' MITEs occurrences**

A region is defined as a genomic sequence with a high density of repeats within a sliding window of 500 bp. Within each region, the number of M1 and M2 repeats was counted. The clusters containing divergent M1 and M2 sequences were included in these results.

| | | *Pithovirus sibericum* | | *Pithovirus mammoth* | | *Pithovirus massiliensis* | |
|---|---|---|---|---|---|---|---|
| | | **M1** | **M2** | **M1** | **M2** | **M1** | **M2** |
| **Regions** | **Total** | 110 | 100 | 109 | 100 | 115 | 79 |
| | **M1 or M2** | 10 | 0 | 9 | 0 | 36 | 0 |
| **Per region** | **Min count** | 1 | 1 | 1 | 1 | 1 | 1 |
| | **Max count** | 11 | 12 | 13 | 17 | 13 | 8 |
| | **Mean** | 4.68 | 3.71 | 4.58 | 4 | 5.05 | 3.01 |
| | **Sd** | 2.12 | 2.26 | 2.31 | 3.06 | 2.98 | 1.64 |

### Table S4. PebbleScout alignments of M1 or M2 against metagenomic reads

The 10 datasets (in bold) with the most reads matching M1 or M2 were assembled and checked for *Pithoviridae* using the divergent MCP (pv_460) as bait.

| SRA ID | PebleScout score | BioSample | Total # reads | M1 reads (BLASTN *E*value<$10^{-10}$) | M2 reads (BLASTN *E*value<$10^{-10}$) | M1+M2 reads per $10^6$ reads | Largest contig w/ M1 or M2 (BLASTN Evalue<$10^{-10}$) | Pithovirus MCP TBLASTN *E*value | Environmental sample |
|---|---|---|---|---|---|---|---|---|---|
| **SRR3989309** | 89,74 | SAMN05421978 | 18336755 | 1211 | 270 | 80,7667442 | 903 | 5,64E-34 | Terrestrial. Deep subsurface. Rock core/Sediment |
| **SRR2090167** | 89,74 | SAMN03842445 | 28636935 | 1725 | 368 | 73,0874306 | 1091 | 1,10E-66 | Groundwater. Rifle well CD01 at 16ft depth; 0.1 micron filter at time point B |
| **SRR11310430** | 84,61 | SAMN14381997 | 6032418 | 317 | 79 | 65,6453183 | 526 | 5,81E-53 | Sediment from asphalt lake |
| **SRR2090164** | 89,74 | SAMN03842442 | 19817017 | 1014 | 217 | 62,1183299 | 886 | 1,18E-23 | Groundwater. Rifle well CD01 at 16ft depth; 0.1 micron filter at time point A |
| **SRR3989312** | 89,74 | SAMN05421984 | 19817017 | 1014 | 217 | 62,1183299 | 675 | 1,18E-23 | Deep subsurface. Groundwater |
| **SRR11310431** | 79,49 | SAMN14381997 | 6170151 | 275 | 63 | 54,7798587 | 762 | 2,26E-43 | Sediment from asphalt lake |
| **SRR5381855** | 79,52 | SAMN06546764 | 1684054 | 72 | 11 | 49,2858305 | 462 | 2,79E-24 | Soil metagenome of an asparagus field |
| **SRR6208705** | 89,74 | SAMN07687476 | 17707616 | 505 | 102 | 34,2790356 | 701 | 1,32E-22 | Terrestrial. Deep subsurface. |
| **SRR5381897** | 74,38 | SAMN06547013 | 3721304 | 90 | 14 | 27,9471927 | 527 | | Soil metagenome of an asparagus field |
| **SRR6211583** | 79,49 | SAMN07687567 | 21572737 | 361 | 61 | 19,5617274 | 496 | 6,22E-25 | Terrestrial. Deep subsurface. |
| ERR2206798 | 74,38 | SAMEA104408696 | 17005406 | 95 | 20 | 6,76255539 | | | Brackish water. Baltic Sea |
| SRR2090165 | 79,49 | SAMN03842443 | 22016389 | 114 | 25 | 6,31347856 | | | Groundwater. Rifle well CD01 at 16ft depth; 0.1 micron filter at time point A |
| SRR3989308 | 79,49 | SAMN05421983 | 22016389 | 114 | 25 | 6,31347856 | | | Deep subsurface. Groundwater |
| SRR1955040 | 89,74 | SAMN03460428 | 84084083 | 296 | 39 | 3,98410719 | | | Soil and sludge samples from the vicinity of pesticide manufacturing unit |
| SRR6212587 | 79,53 | SAMN07687597 | 22970382 | 73 | 11 | 3,65688302 | | | Terrestrial. Deep subsurface. |
| SRR2090170 | 89,74 | SAMN03842448 | 20939869 | 54 | 13 | 3,19963797 | | | Groundwater. Rifle well CD01 at 16ft depth; 0.1 micron filter at time point C |
| SRR3989314 | 89,74 | SAMN05422002 | 20939869 | 54 | 13 | 3,19963797 | | | Terrestrial. Deep subsurface. Rock core/Sediment |
| SRR4388699 | 74,38 | SAMN03842451 | 21838918 | 40 | 9 | 2,2437009 | | | Groundwater. Rifle well CD01 at 16ft depth; 0.1 micron filter at time point D |
| SRR15669522 | 79,49 | SAMN21040050 | 51610118 | 94 | 17 | 2,15074106 | | | Groundwater. |
| SRR3546452 | 89,74 | SAMN04999992 | 53343210 | 84 | 22 | 1,98713201 | | | Deep subsurface groundwater filtered through 0.2 um and collected on 10 kDa filter |
| SRR10912807 | 74,39 | SAMN13674977 | 52979393 | 82 | 11 | 1,7553995 | | | Seawater. Antartica |
| SRR3724388 | 74,38 | SAMN05224416 | 36371555 | 44 | 11 | 1,51217071 | | | Marine. Intertidal zone |
| SRR8931195 | 74,4 | SAMN11466655 | 30200907 | 23 | 22 | 1,49002148 | | | Groundwater. |
| SRR3725730 | 79,49 | SAMN05224444 | 52357685 | 64 | 13 | 1,47065326 | | | Marine. Intertidal zone |
| SRR10912892 | 74,38 | SAMN13674977 | 59432429 | 62 | 10 | 1,21145982 | | | Seawater. Antartica |
| SRR8893624 | 74,38 | SAMN11412375 | 41695761 | 19 | 9 | 0,67153109 | | | Waste water |
| SRR10912798 | 79,54 | SAMN13674978 | 56855335 | 25 | 6 | 0,54524347 | | | Seawater |
| SRR636569 | 84,63 | SAMN01828240 | 3,42E+08 | 77 | 21 | 0,28660667 | | | Waste water |

### Table S5. HOGs related to transposase or integrase

| | | | | In cluster with | | **Best Foldseek match to a transposase or integrase (probability > 0.5)** | | |
|---|---|---|---|---|---|---|---|---|
| | HOG | Gene | Size (aa) | M1 | M2 | Annotation | Probability | E-value |
| *Pithovirus massiliensis* | HOG248 | pmas_554 | 65 | x | x | AF-X8F9W3-F1 Mutator family transposase | 0.887 | 2.46E-01 |
| | | pmas_124 | 51 | x | x | AF-A0A133CJD6-F1 Site-specific integrase | 0.692 | 1.87E+00 |
| | | pmas_125 | 72 | x | x | | | |
| | | pmas_355 | 54 | x | x | | | |
| | | pmas_397 | 56 | x | x | AF-A0A1D6J6D6-F1 DUF659 domain-containing protein (Transposase-like protein with no known function) | 0.992 | 4.09E-01 |
| | | pmas_398 | 72 | x | x | | | |
| | | pmas_490 | 112 | x | | | | |
| | | pmas_491 | 54 | x | | | | |
| | | pmas_552 | 65 | x | x | | | |
| | | pmas_67 | 56 | x | | | | |
| | | pmas_83 | 60 | x | x | | | |
| *Pithovirus sibericum* | HOG567 | ps_41 | 52 | x | x | | | |
| | | pv_143 | 85 | x | x | | | |
| | | ps_381 | 52 | x | x | | | |
| *Pithovirus mammoth* | | pmam_133 | 52 | x | x | | | |
| | | pmam_241 | 50 | x | x | AF-A0A133CJD6-F1 Site-specific integrase | 0.663 | 1.97E+00 |
| *Pithovirus massiliensis* | HOG272 | pmas_168 | 68 | x | | | | |
| | | pmas_212 | 103 | x | x | AF-E9Q492-F1 PiggyBac transposable element-derived 1 | 0.692 | 7.22E-01 |
| | | pmas_352 | 74 | x | | | | |
| | | pmas_425 | 52 | x | x | | | |
| *Pithovirus sibericum* | | pv_335 | 90 | x | x | | | |
| *Pithovirus mammoth* | | pmam_295 | 56 | x | x | | | |
| | | pmam_395 | 55 | x | x | | | |
| | | pmam_445 | 90 | x | x | | | |

### Table S6. Functional annotation of genes from *Pithovirus sibericum* based on Alphafold prediction and Foldseek alignments

| Gene | Previous annotation | Alphafold/Foldseek annotation | Within in repeat-rich region | Is HGT |
|---|---|---|---|---|
| pv_3 | Uncharacterized protein | Ubiquitin thioesterase OTU1 | 0 | 0 |
| pv_4 | Uncharacterized protein | Proliferating cell nuclear antigen | 0 | 0 |
| pv_9 | Uncharacterized protein | redox-related protein | 0 | 0 |
| pv_38 | Conserved protein | Kinase | 0 | 0 |
| pv_39 | Conserved protein, partial | Protein kinase | 0 | 0 |
| pv_50 | Uncharacterized protein | Serine/threonine-protein kinase Chk1 | 0 | 0 |
| pv_51 | Uncharacterized protein | TATA box-binding protein-like | 0 | 0 |
| pv_66 | PolyA pol reg subunit | Cap-specific mRNA (nucleoside-2'-O-)-methyltransferase | 1 | 0 |
| pv_95 | Uncharacterized protein | Ricin B-like lectin | 0 | 0 |
| pv_101 | Uncharacterized protein | Kinase | 0 | 0 |
| pv_102 | Uncharacterized protein | Protein kinase | 1 | 0 |
| pv_105 | Uncharacterized protein | ELAV-like protein | 0 | 0 |
| pv_109 | Uncharacterized protein | Metacaspase | 0 | 0 |
| pv_110 | Uncharacterized protein | Ras-related protein Rab-6 | 0 | 0 |
| pv_112 | GTP binding protein | Ras-related GTP-binding protein | 0 | 0 |
| pv_115 | Uncharacterized protein | ATP-dependent RNA helicase | 0 | 0 |
| pv_135 | Uncharacterized protein | Phosphomevalonate kinase | 0 | 0 |
| pv_141 | Uncharacterized protein | Acyltransferase | 0 | 0 |
| pv_144 | Uncharacterized protein | Acetyltransferase | 1 | 0 |
| pv_145 | Uncharacterized protein | Protein kinase | 0 | 0 |
| pv_159 | DNA repair exonuclease | Nuclease SbcCD subunit D | 0 | 0 |
| pv_166 | Uncharacterized protein | Putative NAD(+)--arginine ADP-ribosyltransferase | 0 | 0 |
| pv_189 | Uncharacterized protein | GTP-binding nuclear protein Ran | 1 | 1 |
| pv_288 | Uncharacterized protein | GTP-binding protein | 1 | 0 |
| pv_290 | Uncharacterized protein | Ras-related protein Rab | 1 | 0 |
| pv_324 | Conserved protein | Exonuclease | 0 | 1 |
| pv_341 | 2OG-Fe(II) oxygenase | Alpha-ketoglutarate-dependent dioxygenase alkB homolog | 0 | 0 |
| pv_352 | Conserved protein | RNA methyltransferase | 0 | 0 |
| pv_379 | Conserved protein | fatty acid-binding protein | 1 | 0 |
| pv_403 | Uncharacterized protein | Nudix hydrolase | 0 | 0 |
| pv_406 | Glycosyltransferase family 2 | Mannan polymerase complex subunit | 0 | 0 |
| pv_408 | Uncharacterized protein | Protein kinase | 0 | 0 |
| pv_423 | Uncharacterized protein | Regulatory subunit of aspartate kinase | 0 | 0 |
| pv_424 | Uncharacterized protein | Protein kinase | 0 | 0 |
| pv_444 | Poly A pol reg subunit | Cap-specific mRNA (nucleoside-2'-O-)-methyltransferase | 0 | 0 |
| pv_445 | Uncharacterized protein | Crossover junction endodeoxyribonuclease RuvC-like | 1 | 0 |
| pv_468 | Uncharacterized protein | F-box containing-protein | 1 | 0 |

**Table S7. Genomic rearrangements and mutations between *Pithovirus sibericum* and *Pithovirus mammoth***

| | Rearrangements types | | | | | | | Total orthologous pairs with rearrangement events | Conserved orthologous pairs without rearrangement event |
|---|---|---|---|---|---|---|---|---|---|
| Repeats regions | Insertions/ deletions | Single nucleotide insertions/ deletion | Substitutions | Inversions | Duplications in *Pithovirus sibericum* | Duplication in *Pithovirus mammoth* | Complex events | | |
| Within | 9 | 5 | 2 | 5 | 1 | 4 | 2 | 228 | 109 |
| Outside | 5 | 2 | 1 | 1 | 1 | 3 | 0 | 13 | 41 |
| | | | | | | | | Chi$^2$ *P*value = $5.5 \times 10^{-7}$ | |
